# Task-driven neural network models predict neural dynamics of proprioception

**DOI:** 10.1101/2023.06.15.545147

**Authors:** Alessandro Marin Vargas, Axel Bisi, Alberto Chiappa, Chris Versteeg, Lee Miller, Alexander Mathis

## Abstract

Proprioception tells the brain the state of the body based on distributed sensors in the body. However, the principles that govern proprioceptive processing from those distributed sensors are poorly understood. Here, we employ a task-driven neural network modeling approach to investigate the neural code of proprioceptive neurons in both cuneate nucleus (CN) and somatosensory cortex area 2 (S1). We simulated muscle spindle signals through musculoskeletal modeling and generated a large-scale, naturalistic movement repertoire to train thousands of neural network models on 16 behavioral tasks, each reflecting a hypothesis about the neural computations of the ascending proprioceptive pathway. We found that the network’s internal representations developed through task-optimization generalize from synthetic data to predict single-trial neural activity in CN and S1 of primates performing center-out reaching. Task-driven models outperform linear encoding models and data-driven models. Behavioral tasks, which aim to predict the limb position and velocity were the best to predict the neural activity in both areas. Architectures that are better at solving the tasks are also better at predicting the neural data. Last, since task-optimization develops representations that better predict neural activity during active but not passively generated movements, we hypothesize that neural activity in CN and S1 is top-down modulated during goal-directed movements.

## Introduction

A central tenet of neuroscience is that neural circuits are shaped by evolution and development to enable ethologically important behaviors. Task-driven modeling constitutes a framework that creates candidate models of the brain by behavioral optimization (1–4), and has emerged as a powerful approach for studying sensory systems (5–9). Task-driven models best predict visually-driven responses (5, 9, 10), and provide insights about the organization and principles of sensory processing (5, 6, 9, 11–13) which can be used to guide new experiments (7, 14). Task-driven approaches have also been developed to modeling the proprioceptive pathway, and were used to study emerging coding properties (15, 16). However, these models have not been used to predict neural activity in the proprioceptive pathway yet.

Proprioception informs the brain about the position, movement and associated forces of the limbs (17–20). Muscle spindles, mechanoreceptors distributed in muscles, constitute the main source of proprioceptive information to the brain. These receptors are sensitive to muscle stretch and stretch velocity, and possibly other muscle properties such as force (21–25). Muscle spindle afferents convey muscle kinematics information through the dorsal root ganglia, the dorsal column nuclei in the brainstem, including the cuneate nucleus (CN) for upper limb muscles, thalamus, primary (S1) and secondary somatosensory cortices, in this order forming the ascending proprioceptive pathway (26–33). Proprioceptive neurons typically show distinct gain modulation profiles during active voluntary movement versus passive limb displacement (31, 34–36). It is currently not well understood how proprioceptive information is transformed along this pathway.

The proprioceptive pathway supports a diversity of sensory and motor-related behaviors (18, 19). It has been hypothesized that proprioceptive signals support motor control and learning (18, 37, 38), localizing the position, velocity and acceleration of body parts in various coordinate frameworks (17) and action recognition (15, 19), i.e. recognizing location-invariant movements. Furthermore, like for other sensory modalities, proprioceptive signals might be transformed to represent sensory information in an efficient way, as proposed by Horace Barlow (39), either by reducing the correlation between sensory inputs, or by enabling faithful reconstruction of sensory inputs (efficient coding hypothesis (40)).

Here, we developed a normative framework to evaluate different hypotheses using task-driven neural network models of the proprioceptive pathway. Expanding upon our prior work (15), we generated a large-scale, synthetic proprioceptive inputs resulting from passive, naturalistic movement, which we used to train task-driven neural network models on 16 tasks. For model training, we did not include any muscle force or activity in the synthetic data but focused only on passively generated movements. We then tested if the network’s learned internal representations resembled those of proprioceptive brain areas, by using these network representations to predict the neural activity in the primate cuneate nucleus (CN) and primary somatosensory cortex (S1 area 2). For this purpose we used experimental data collected from nonhuman primates (NHP) during both active and passive center-out arm movements (36, 41) (Figure 1). First, we show that our task-driven models could accurately predict single-trial neural dynamics in both conditions, outperforming classical encoding models. Second, we show that the choice of behavioral tasks and neural network architectures are crucial to the development of more “brain-like” representations. In particular, tasks which predict the kinematics (position and velocity) of the limb, irrespective of the choice of body reference system, were the best to predict neural activity of proprioceptive neurons in both CN and S1. This suggests that a motor-efference copy might be sent to CN and S1 modulating their activity. We show that for the majority of tasks, architectures that have better task performance, are also better at explaining the neural data. Finally, task-trained models outperform untrained models during voluntary movement, but not during passive movements. This suggests that there might be a top-down modulation of CN and S1 activity during goal-directed movements, while such modulation is likely absent during passive movements.

**Figure. 1:**
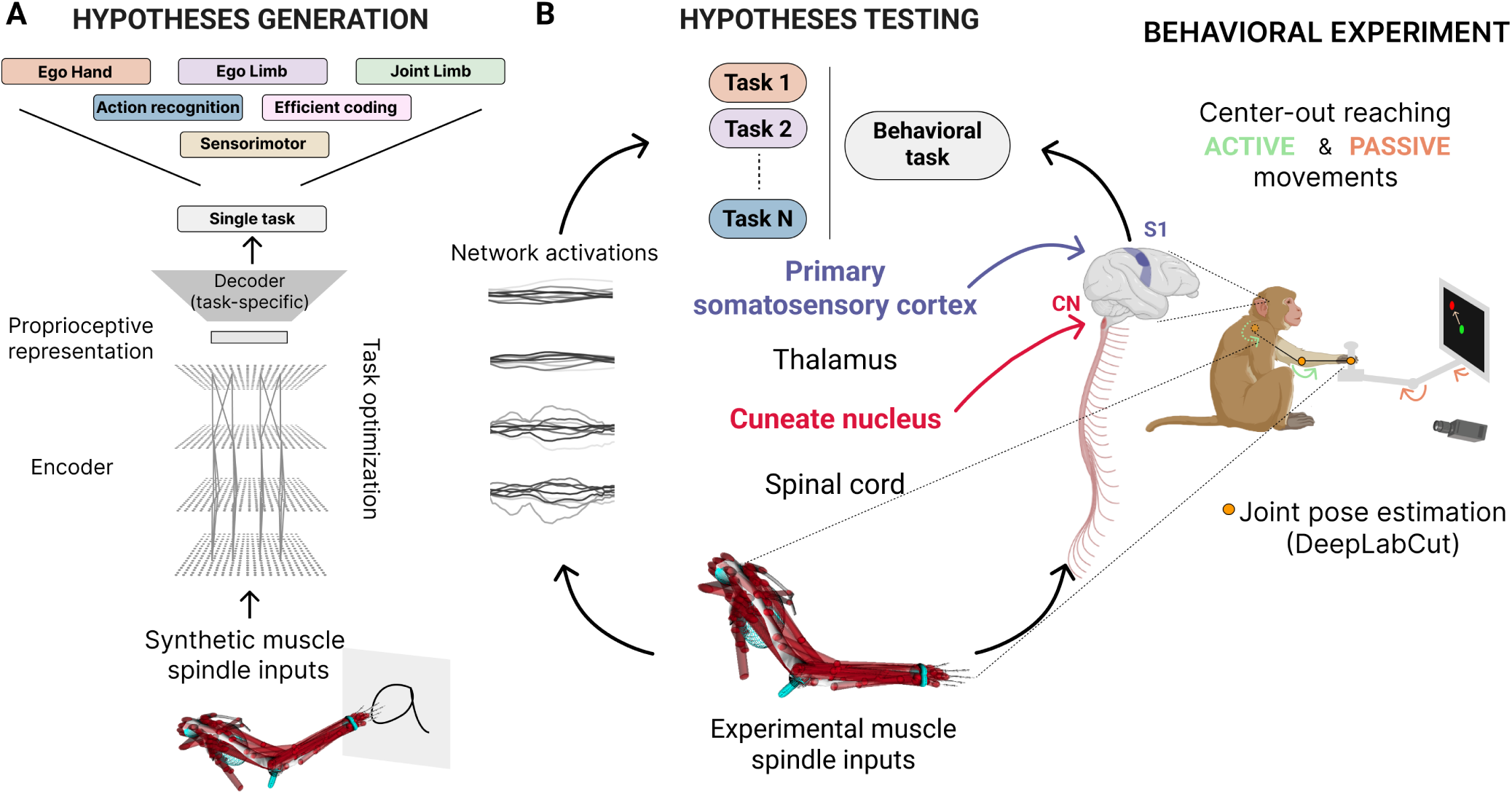
Task-driven models as a normative framework for interrogating proprioception. **A.** We created a normative framework to study the neural code of proprioception. Using synthetic muscle spindle inputs derived from musculoskeletal modeling, we optimized deep neural networks to solve different computational tasks in order to test hypotheses (N=16 tasks). Task-driven models give rise to learned representations across the artificial neural network models. **B.** We interrogated which type of hypothesis creates models that best generalize to explain the activity of neurons in the cuneate nucleus (CN) of the brainstem and in the primary somatosensory cortex (S1, area 2) of non-human primates performing a center-out reaching task comprising active and passive movements.

## Results

### Task-driven models as a normative framework for interrogating proprioception

We developed a normative framework to investigate the neural code of proprioception. The proprioceptive system receives distributed information from muscle spindles and it is an open question how it integrates this information to subserve behavior. We considered 16 different proprioception-based tasks that the proprioceptive system might be optimized for. By employing a task-driven modeling approach, we trained deep neural network models on those tasks and investigated which of those hypotheses force the network to develop representations that best resemble primate proprioceptive representations. To tackle this question, we simulated large-scale synthetic muscle spindle receptor inputs during naturalistic movement and created hierarchical deep neural network (DNNs) models that were trained to solve these tasks directly from receptor inputs, akin to what the biological system has to do. We evaluated whether the task-driven neural network models could predict neural data recorded in five non-human primates (NHP) performing a center-out reaching task (Figure 1).

### Building task-driven models of proprioception that solve ethologically relevant tasks

The proprioceptive pathway supports a myriad of sensory and motor-related behaviors (18, 19, 42). For our normative framework, we tested 16 different hypotheses using different neural network architectures (Figure 1). Artificial neural networks comprise multiple layers of simplified units (“neurons”) whose connectivity patterns mimic the hierarchical, integrative properties of anatomical pathways (1, 2). Like Sandbrink et al. 15, we considered both temporal convolutional networks (TCNs), and a recurrent neural networks (long short-term memory (LSTM)), which impose different inductive priors on the computations (See methods).

We tested 16 different hypotheses (Figure 2C and Table 4). Specifically, we implemented the “EgoHand” hypothesis, which predicts separately or in combination the position, velocity and acceleration of the end-effector in egocentric coordinates from the distributed muscle spindle input (N = 4 tasks - “EgoHand hypothesis” column of Figure 2C). Similarly, we considered the “EgoLimb” hypothesis that also includes the kinematics of the elbow as prediction target in addition to the end-effector (4 tasks). We developed four tasks to predict joint kinematics (“JointLimb”). All these supervised tasks predict the state of the body in various reference frames, which is consistent with prior proposals on the role of proprioception (19, 20, 42). We included the action recognition hypothesis (15), where networks were trained to classify from the proprioceptive input the movement type performed in different workspace locations. We implemented a sensorimotor hypothesis (18), where the model had to predict the torques needed to reach a target provided by a separate input (cue) from the current state. To be able to use the same encoders and inputs as for the other tasks, we first trained a reinforcement learning agent to control a 4 DoF arm to reach targets in the 3D space (Supp. Fig. 1A,B). Through a teacher-student approach, we trained the same encoders to regress learned joint torques of the agent from the muscle spindle inputs and a cued target input. The student network could accurately reach the different targets (See methods, Supp. Fig. 1C,D). Furthermore, we developed two unsupervised tasks: redundancy reduction and autoencoders which reconstruct the proprioceptive inputs. For the former, we developed a modified version of the Barlow Twins self-supervised loss, initially proposed for visual processing (43). The latter aimed to develop networks that can robustly reconstruct the proprioceptive inputs. Those tasks represent variants of the efficient coding hypothesis according to which sensory systems have evolved to compress the input statistics in a meaningful way (39, 40). We emphasize that we used the same sensory statistics and architectures for all tasks, but only varied the decoders, which are task-specific (Figure 1A). Overall, we trained 350 models architectures on each of those 16 tasks corresponding to a total of 5600 neural network models (See methods).

**Figure. 2:**
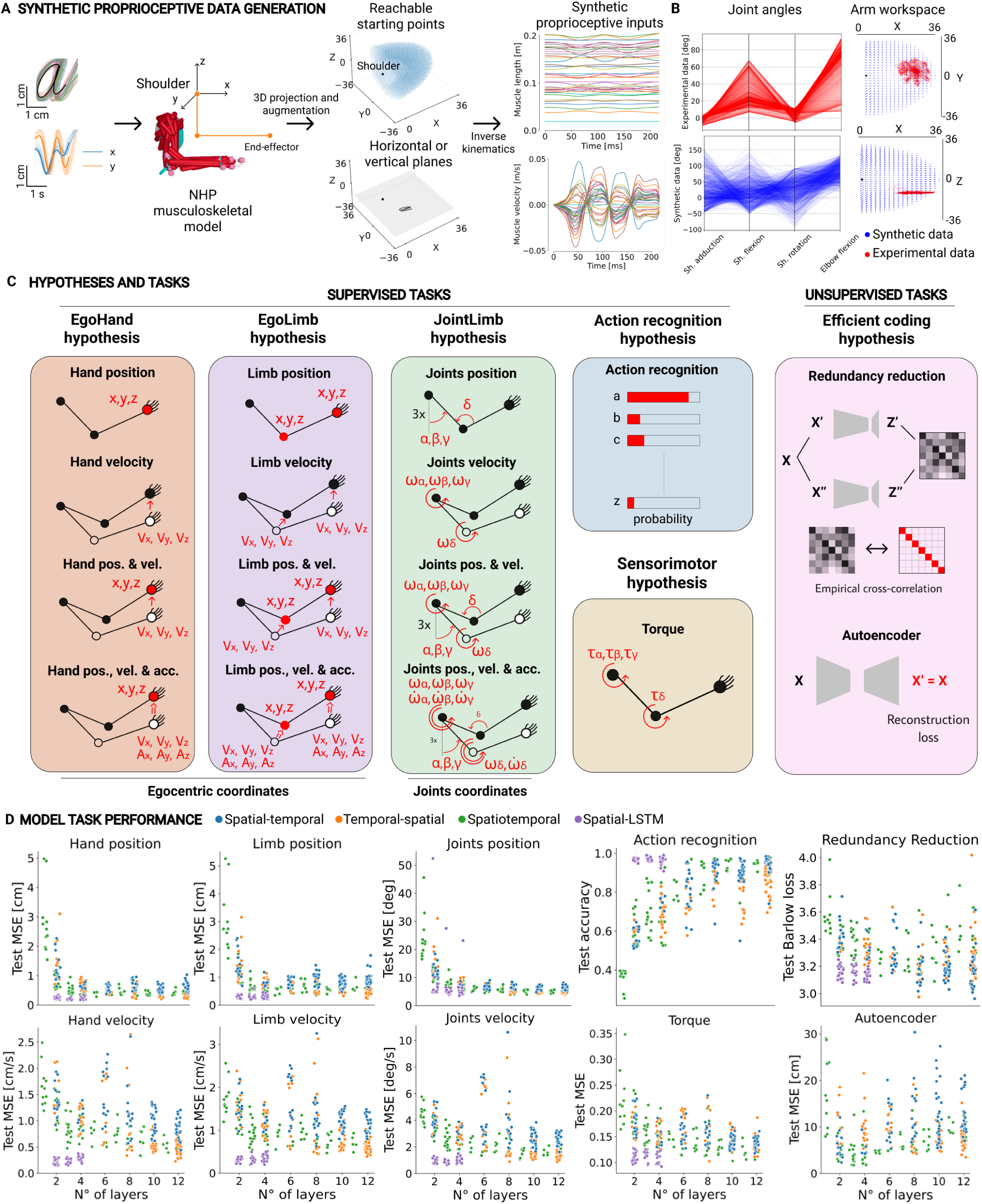
Building task-driven models of proprioception that solve ethologically relevant tasks. **A.** Diagram of the computational pipeline to compute realistic synthetic proprioceptive inputs from 3D character trajectories. In brief, 2D character trajectories are augmented and projected into 3D space using a 2-link 4 degree of freedom (DoF) arm model by randomly selecting candidate starting points in the workspace of the NHP arm. From those movements, muscle lengths and velocities are computed using inverse kinematics and musculoskeletal modeling. **B.** Left: Distribution of joint angles and trajectories in the behavioral (top) and synthetic (bottom) data, illustrating that the movement statistics in the synthetic dataset are designed to (broadly) encompass the biological movements of NHPs during the center-out reaching task. Right: workspace trajectory starting points of the synthetic dataset (blue), encompassing the behavioral workspace (red). Note that experimental center-out trajectories themselves are not part of the synthetic dataset. **C.** In total, 16 computational tasks were designed to reflect different hypotheses about functional proprioceptive processing. Each hypothesis contains one or several objectives, grouped by similarity. The background of the panel is color coded based on the hypothesis and it will be used throughout the manuscript. For each task, the learning objective is highlighted in red with each arm pictogram. **D.** Test performance of each network model, on selected tasks (N = 350 models except the autoencoder task where N=295), with respect to the number of layers (i.e. model depth). For the regression tasks, we used the mean squared error (MSE), for the action recognition task, the classification accuracy, for the autoencoder task, the relative error for the muscle length and for the redundancy reduction task, the Barlow loss (See methods). Four types of deep neural network architectures were designed to integrate proprioceptive signals in different ways: spatial-temporal, temporal-spatial, and spatiotemporal TCNs, and spatial-LSTM. The color code of each point reflects the architecture type.

We assessed the performance of the trained neural networks using task-specific metrics to ask whether the designed models could solve all these different tasks (Figure 2E and Supp. Fig. 2). For each task, we observed that neural networks can achieve strong performance and that deeper neural network models correlate with better task performance. For instance, models could predict the location of the end-effector or the elbow with a 1cm accuracy and they could achieve higher than 99% action recognition classification accuracy. Additionally, they could also reconstruct the muscle state with only 0.01% relative muscle length error for autoencoders (Figure 2E). However, neural networks trained on tasks that included acceleration in the regression target showed poorer target prediction ability for other variables (position and velocity) compared to the networks that were trained to specifically predict only those variables (Supp. Fig. 2). Due to their recurrence, the LSTM networks achieved strong performance with fewer encoding layers.

In summary, we developed a normative framework to test hypotheses describing how behavioral or computational constraints might shape proprioceptive processing. We generated a naturalistic, large-scale dataset of proprioceptive signals and used it to train thousands of neural network models to successfully perform various tasks.

### Reaching data to test task-driven models

While the task-driven models were trained on synthetic data only, we wanted to assess whether their internal representations developed through task-optimization resembled those of proprioceptive areas. To this end, we tested the generalization ability of these task-driven models from synthetic data to biological data by predicting neural activity recorded in non-human primates (rhesus macaques) during a center-out reaching paradigm (36, 41). The center-out behavioral paradigm contains both active and passive upper limb movements corresponding to distinct proprioceptive conditions. In the active condition, the NHP controlled a manipulandum to reach equidistant targets within a circle on a screen whereas, in the (randomly interspersed) passive condition, the robotic manipulandum applied an unexpected force perturbation to the NHP arm towards one of the target directions for a duration of 130 ms (Figure 3A). Neural activity in cuneate nucleus (CN) and area 2 of somatosensory cortex (S1) was recorded with Utah arrays in NHPs performing the task at expert level. We used rewarded trials from three recording sessions per area, from each of NHPs, each including between 10 and 93 neurons (total CN: 159; total S1: 87) and varying number of trials (Figure 3B). Data from one NHP contained simultaneous recordings in CN and S1, labeled throughout the text as L and S1L respectively. Examples of single-trial neural activity are shown for both active and passive conditions for one example CN unit, each with direction-dependent firing rates (Figure 3C, E).

**Figure. 3:**
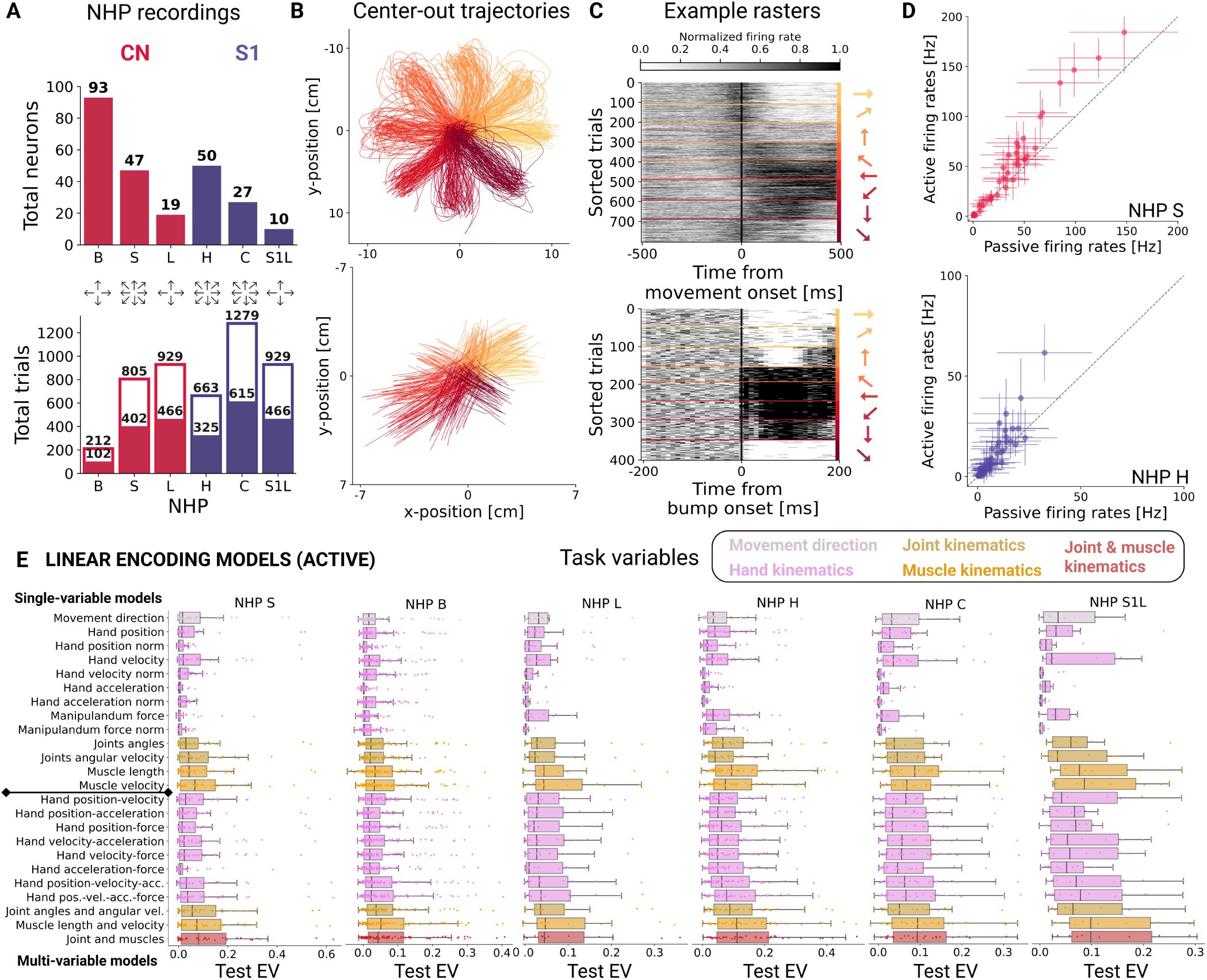
Center-out reaching experiments, and linear encoding models of neural activity. **A.** Size of experimental datasets. Top: total number of neurons per NHP per area. Bottom: total number of trials for active (hollow bars) and passive (full bars) conditions per NHP. Note that individuals L and S1L are the same NHP hence the same number of trials. The color code denotes the area, CN (red) and S1 (blue). The number of different reach directions are indicated for each NHP (4 or 8). **B.** Planar x-y coordinates of the screen cursor representing end-effector trajectories of the NHP during the center-out reaching task for one example behavioral session during active reaches (top, N=805 trials) and passive perturbations (bottom, N=402 trials). The color code indicates the target direction. **C.** Raster plot of the normalized firing rate, for one example CN neuron, aligned to the movement onset (vertical black line) and sorted by movement direction for one session, in active (top) and passive trials (bottom). **D.** Pairwise single-neuron comparison of trial-averaged firing in the active and passive conditions for NHP S (CN; N=47 neurons, top) and NHP H (S1; N=50 neurons, bottom). Vertical and horizontal bars represent the SEM across trials. **E.** Distributions of the performance across single neurons of the baseline linear encoding models using task and behavioral variables, alone (single-variable models) or in combination (multi-variable), for all NHPs. The color code reflects the group of the behavioral variable. Note, the muscle kinematics (muscle length and velocity; orange) is the input to our neural network models.

To predict center-out neural activity using task-driven models, we inferred the muscle spindle inputs for our models using DeepLabCut (36, 44) to estimate the location of limb joints during the center-out reaching task. We extracted muscle kinematics via musculoskeletal modeling and inverse kinematics (See methods, Supp. Fig. 3 A, B); the latter step was done in the same manner as for the synthetic dataset. In this way, we can use center-out experimental data as novel test input data to our neural network models in order to compare artificial to biological representations.

### Baseline linear encoding models

Before predicting neural activity via task-driven neural network models, we first looked at how much variance of the neural activity could be captured using classical linear encoding models (45). The activity of proprioceptive neurons is known to be correlated to both kinematic and dynamic variables (31, 34, 36, 41, 45). Therefore, we fitted single-trial neuron activity through ridge regression using as input a variety of movement-related variables (single-variable models), as well as their combinations (multi-variable models) (Figure 3F, Supp. Fig. 3D). We will refer to these models as *linear encoding models* throughout the manuscript and they define the baseline to which we will compare task-driven models. The variables included in the linear models are hand kinematics (position, velocity, acceleration), joint kinematics (angular position and velocity) and muscle kinematics (length, velocity). We also included a two-dimensional interaction force between the NHP’s hand and the grasped manipulandum. Moreover, we also considered the magnitude of the hand kinematic vectors and of the manipulandum force vector as a regressor. The experimental trials were split in 80% train and 20% test set and we cross-validated the ridge regularization hyperparameter based on the train trials only. We note that ridge regression was more robust than generalized linear models (GLMs) with Poisson distribution (See methods, Supp. Fig. 3C). We quantified the accuracy of the predictions by computing the explained variance (EV) on test session trials for each neuron (See methods).

Consistently with previous works (41, 45, 46), we observed that models fitted using variables spanning the whole-arm kinematics - i.e., multi-joint and multi-muscle kinematics - provided better prediction of neural responses in CN and S1, in both active and passive movement conditions, than did hand kinematics only (Figure 3F, Supp. Fig. 3D). Additionally, we observed that multi-muscle kinematics achieve higher explained variance across all multi-variable models for both CN and S1. This result suggests that CN and S1 neurons are best tuned to muscle kinematics, rather than hand or joint kinematics.

### Task-driven models predict neural dynamics

The central question of this study is whether task-driven models trained only on synthetic data can develop representations that generalize to predict neural dynamics of the primates proprioceptive system. We emphasize that task-driven models have not been trained on a center-out reaching task, with the exception of the torque regression task. Rather, network models were trained with different movement statistics in order to test generalization from synthetic to real movement data and from one task to another (the center-out reaching task).

In contrast to previous works on other task-driven models of sensory systems, where the predicted neural activity was typically trial-averaged (e.g., (5, 6, 9, 11, 13) but see (12)), here we aim to predict the temporally varying firing rates of CN and S1 neuron for each center-out reaching movement using the inferred muscle spindle inputs (See methods). To this end, we froze the neural network models after training on synthetic data. Then, we retrieved the resulting internal representations at each model layer using as input the simulated proprioceptive inputs during the center-out limb movements. Single-neuron activity was predicted using as regressors the 75 first principal components (PCs) of the network’s internal representations. We kept the number of PCs fixed along the network hierarchy to keep the same number of dimensions irrespectively of the layer size. The number of PCs was crossvalidated (Supp. Fig. 4A) and it is lower than the number of proprioceptive inputs (i.e. muscle length and velocity of 39 muscles; 2 *×* 39 = 78). Just as for linear models, we used ridge regression to predict the neural activity from the PCs of each layer. As a first sanity check, we found that the distributions of explained variance were similar between training and testing trial datasets across all NHPs, which suggests that there is no overfitting (Supp. Fig. 4C).

We want to illustrate initial results focusing on one example task-driven model: a spatial-temporal model with 12 layers trained on the hand position and velocity task (EgoHand hypothesis). The task-driven neural network model could predict single-neuron activity and capture the single-trial dynamics (Figure 4A). We quantified the encoding variability of task-driven models by analyzing the distribution of explained variance both across neurons, NHPs and experimental conditions (active vs. passive). We found a wide range of EV across different neurons for both active and passive movement conditions (Figure 4B). Qualitatively, we observed that task-driven predictions for some neurons capture part of the neural dynamics that linear models cannot (Supp. Fig. 5). Indeed, predictions obtained from task-driven models explain the neural activity significantly better than linear models for almost all neurons and for both active and passive conditions (Figure 4C). This holds true for all NHPs and is statistically significant when deep layers of the task-driven models are used to predict the neural activity (Supp. Fig. 4 D,E).

**Figure. 4:**
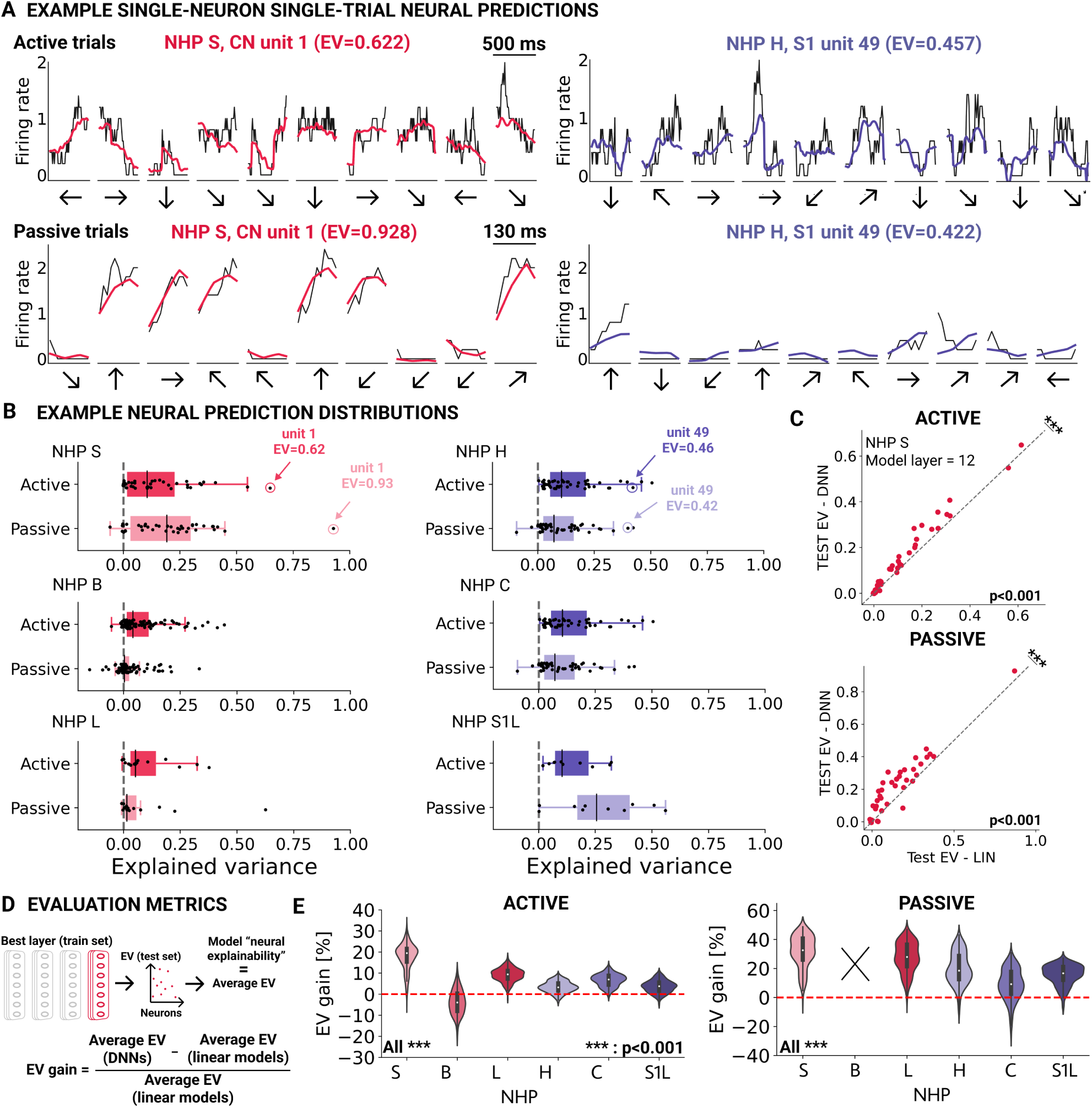
Task-driven neural network models predict neural dynamics during active and passive movements. **A.** Example of time-varying neural predictions (on test trials) for active (top) and passive (bottom) movements, for one example unit in CN (left) and one in S1 (right). The black lines corresponds to ground truth spike firing rates (binned in 50 ms windows) while the colored lines correspond to the model predicted rates. The movement direction of each trial is indicated by an arrow on the x-axis. These example predictions are shown using the best-predicting layer of a 12-layer spatial-temporal neural network model trained on the hand position and velocity task. **B.** Example distributions of single-neuron neural explainability for the best model layer (using the same model as panel A) across all neurons for all NHPs and for active and passive movements. Each neuron is represented by a black dot and the neurons shown in panel A are highlighted. **C.** Pairwise comparison of single-neuron explained variance (NHP S; CN, N=47 neurons) between the last layer of one example deep neural network model shown in B and the baseline linear encoding model (as in Figure 3) using muscle spindle inputs (muscle length and velocity). The statistical significance was computed with the Wilcoxon signed-rank test. **D.** Schematic illustration of the model “neural explainability” metric and the computation of the gain in explained variance between linear models and deep neural network models (DNNs). **E.** Distributions of EV gain for 350 models trained on the hand position and velocity task, for all NHPs in the active (left) and passive (right) conditions. We computed the statistical significance using a two-sided paired t-test between active and passive conditions for each NHP. The red line indicates DNN performance equal to that of baseline linear models.

In order to facilitate comparisons between different models, we defined a metric called “neural explainability” (EV). This metric represents the average test explained variance across neurons per each NHP for the optimal predicting layer, which is selected from the training set (Figure 4D). For linear models, this value is simply the average explained variance across neurons. In this way, we can assess whether task-driven models are able to develop representations that better predict the neural activity than linear models for a given task. Therefore, we compared the neural explainability of task-driven networks trained on the hand localization and movement task to linear models (Figure 4E and Supp. Fig. 5). Our analysis revealed that the majority of task-driven networks provided better neural predictions for all NHPs. This suggests that task optimization can lead to the development of robust representations across various network architectures, resulting in a maximum gain of up to 40% for the active condition and up to 50% for the passive condition in EV per NHP.

For each task, we trained different types of network architecture to ensure robustness in the developed representations and to investigate which architecture types gave better representation, particularly regarding specific types of integration. We evaluated the performance of the network architecture by comparing the EV per NHP, regardless of the task used for training. We found that TCNs performed significantly better than spatial-LSTMs in terms of variance explained for both active and passive condition (Supp. Fig. 6A,B), leading us to focus on the 300 TCNs for each task. Among the TCN types, we found that spatiotemporal networks were better at predicting S1 neural activity during active conditions, suggesting a possible simultaneous muscle-temporal integration during goal-directed movements in S1. Overall, our findings suggest that task-driven models generalize to predict proprioception-related neural dynamics, outperforming classical encoding models.

### Task-driven outperforms data-driven models

We asked if the ability to predict neural activity from neural network models was simply a result of their higher capacity (for fitting data). Thus, we tested whether the same neural network architectures directly trained to predict the neural activity could achieve similar or better performance than task-driven models. Therefore, we trained the same network architectures to regress the firing rate during active movements end-to-end and performed a pairwise comparison to the corresponding task-driven performance. We found that task-driven networks strongly outperform datadriven networks, with the latter exhibiting poor neural predictions (Supp. Fig. 6C). These results provide support for the use of task-driven models to predict neural activity, which involve the use of large-scale (even synthetic) datasets to train over-parameterized neural networks. On the contrary, models trained with the data-driven approach usually require a large amount of experimental data.

### Task-driven outperform untrained models

Starting from the same random, untrained initialized models, we trained each model on different tasks. We reasoned that if task-optimization drives the emergence of proprioceptive-like representations in neural networks, task-trained models should yield better predictions than the corresponding random, untrained models (UNT) and than baseline linear models.

Consequently, we compared neural predictions obtained from the ten best models trained on different tasks to predictions from untrained models, and also from linear models fitted on muscle (Linear - muscles) and hand-only (Linear - hands) kinematics (Figure 5A). First, we observed that for almost all primates, both trained and untrained network predictions are better than linear model predictions, in both active and passive conditions. Second, task-driven models yield a wide range of explained variance: some tasks yield representations that explain the neural activity better than corresponding untrained models highlighting the benefit of task optimization, whereas some tasks surprisingly yield representations worse than the corresponding untrained models.

**Figure. 5:**
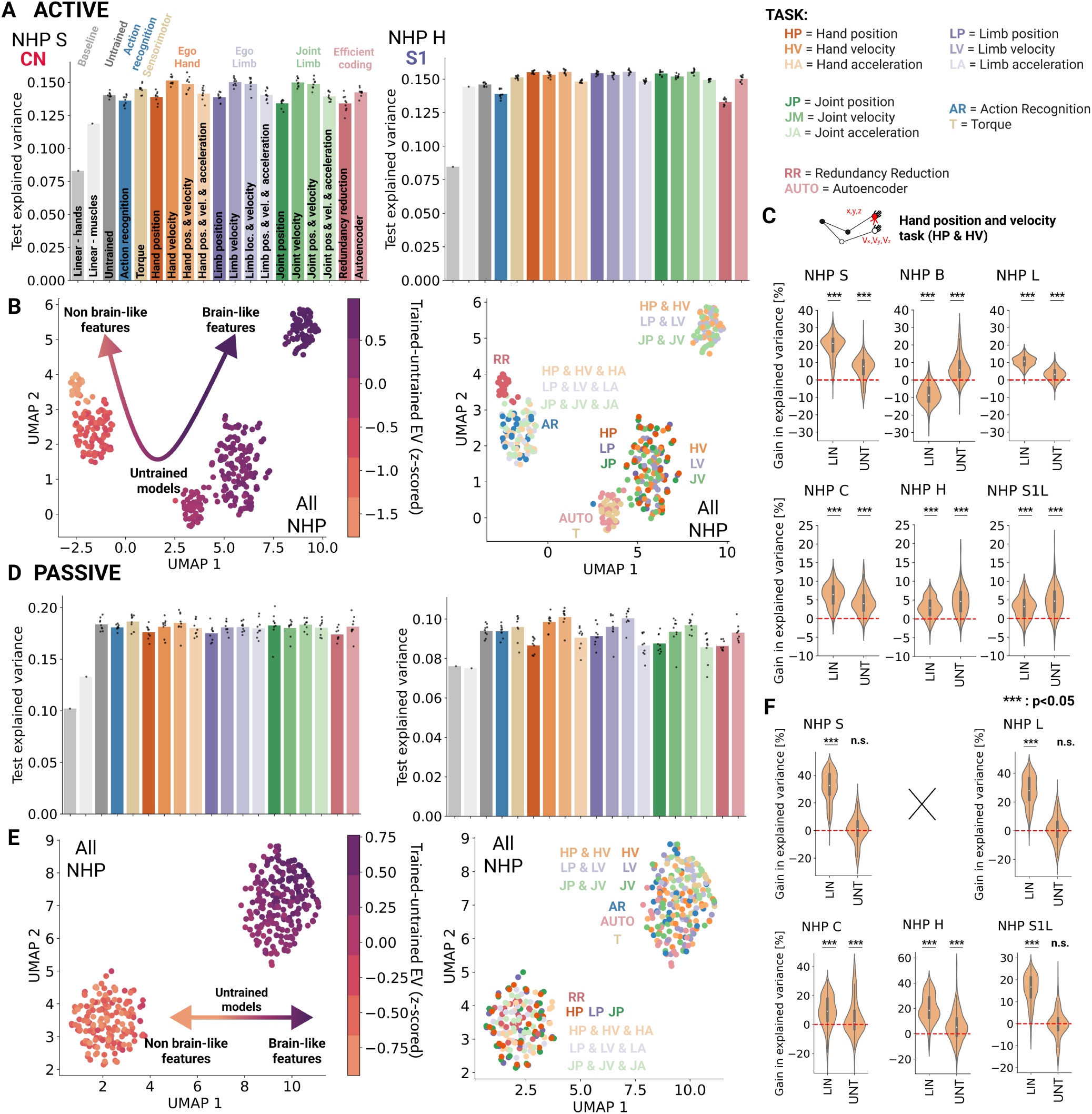
Task comparison show that kinematic-based tasks develop the most “brain-like” features and outperform untrained models. **A.** Distributions of the test neural explainability of selected best 3% model for all tasks and NHP S (CN, left) and NHP H (S1, right), compared with predictions from their randomly initialized counterpart models (dark gray; Untrained) and classical linear models such as hand position and velocity (Linear - hands) and muscle length and velocity (Linear - muscles). The color group code represents the hypothesis to which the task belongs whereas the label for each model task (TASK) is outlined in the task legend. **B.** Low-dimensional UMAP embedding space of the difference in neural explainability between task-driven and untrained networks. Each datapoint represents the z-scored neural explainability associated to a single NHP of a group of networks (N=73) belonging to a given task. Left: we color coded each point based on the corresponding average neural explainability. Right: we color coded each point according to the task identity. **C.** Distribution of the gain, i.e. normalized relative difference, in explained variance when comparing *all* neural network models trained on the hand position and velocity task (TASK: HP & HV) to the linear models (Linear - muscles) and corresponding untrained models (Untrained). The red dashed line indicates models for which there is no difference in explained variance. We note that comparison to linear is repeated from Figure 4E for ease of comparison. The results of two-sided paired t-tests and each random vs. task distribution pair are shown (***: *p <* 0.05). **D.** Same as **A.** but with neural predictions during passive movements during which force perturbations were unexpectedly applied for 130 ms towards one of the equidistant directions. **E.** Same as **B.** but for passive neural predictions. **F.** Same as **C.** but for passive predictions where we do not show the gain in explained variance for NHP B because of the poor performance of all untrained and linear models, which result in exceedingly high gain percentages.

We found that models trained on the hand position and velocity task (HP & HV) were consistently the best across NHPs and movement conditions (Figure 5A and Supp. Fig. 7) . We further tested whether this was true for all models trained on this task or only for specific models that achieved superior neural predictions by comparing, regardless of their task performance, the gain in explained variance relative to the baseline and the corresponding untrained models for all models trained on that task (Figure 5C). We observed that on average task-driven models explain the neural activity significantly better than linear models, reaching a gain up to 40% for the active case and up to 50 % for the passive one.

Interestingly, while task-driven models typically outperformed untrained models significantly under active conditions, this was not always the case for passive conditions (Figure 5D, F) . Nevertheless, this difference could be attributed to the smaller number of experimental trials available during the passive condition as well as the shorter duration of passive perturbations (130ms long). As such, we controlled for these factors by computing neural predictions during active movement that matched the number of passive trials and duration of those movements, i.e. by randomly removing 50% of the active trials and by randomly selecting a sub-interval of 130ms length per trial. Doing so, task-trained models still outperform untrained models in the active condition (Supp. Fig. 6D,E). Overall, these findings corroborate that proprioceptive information might be differently processed during goal-directed and passive, unexpected movements (31, 34–36).

### Kinematic-based tasks explain most variance

We sought to identify which of the 16 tasks drove neural networks to develop more brain-like neural representations by contrasting neural predictions across all the 300 TCN networks and five NHPs. To this end, we wanted to quantify the contribution of each task optimization by comparing the gain in neural predictability that task-driven models can achieve to the corresponding untrained models. To also make the analysis robust with respect to the network architecture, we concatenated the normalized difference of neural explainability between task-trained and untrained models associated to a single NHP for (random) groups of networks belonging to the same task (4 groups of 73 networks per task) and used it as a feature and reduced the dimensionality using UMAP. Therefore, the resulting matrix consist of 4 groups x 6 NHPs x 16 tasks as number of rows and 73 networks (trained – untrained EV) as number of features (See methods).

For the active condition, the networks clustered into five different groups, two that performed better than untrained models, one that was equal and two that were worse (Figure 5C). These clusters also emerged when using PCA, a linear, parameter-free approach (Supp. Fig. 8C). The best cluster in terms of neural predictions only included models simultaneously regressing the position and velocity of the hand and limb (HP & HV, LP & LV, JP & JV tasks), i.e. it contained all tasks that predicted posture and velocity independently of the choice of coordinate framework (i.e. egocentric or joint-angle coordinates). Those models significantly outperformed untrained networks and the other tasks (Wilcoxon signed-rank test; Supp. Fig. 8E).

The second best cluster comprised networks trained to regress either the position or the velocity of either the hand or the limb (HP, LP, JP, HV, LV, JV). The third best cluster contained the autoencoder (AUTO) and the torque regression (T) tasks, but had task-trained neural explainability comparable to that of untrained models. The remaining two clusters had neural explainability significantly worse than untrained models (Wilcoxon signed-rank test, Supp. Fig. 8E,F) and includes tasks aiming to regress the location, velocity and the acceleration of the limb (HP & HV & HA, LP & LV & LA, JP & JV & JA task), recognize actions (AR) and decrease the redundancy within sensory inputs (RR). Among all tasks, we found that redundancy reduction constitutes the task that achieved the poorest neural predictions.

Overall, we found that tasks involving similar regression targets, e.g. predicting only the position or only the velocity of the hand or the limb, are grouped together as they provided similar levels of explained variance (Figure 5C; Wilcoxon signed-rank test, Supp. Fig. 8E). Therefore, we reasoned that these behavioral tasks, as they need to perform similar computation, involve representations that are independent of the target reference frame (i.e., egocentric vs. joint reference frame). Moreover, multi-task training is beneficial to obtain more “brain-like” representations as networks trained to regress both position and velocity outperform networks trained to regress either position or velocity alone. However, when the regression targets also included acceleration, we observed the emergence of “non brain-like” representations and we hypothesize it might be related to the poorer task-performance of these networks compared to the other tasks.

In contrast, for the passive condition, we observed only two main task clusters, each comprising several tasks (Figure 5D; PCA: Supp. Fig. 9C,D; Wilcoxon signed-rank test, Supp. Fig. 9 E)). One cluster contained models whose task-trained neural explainability is slightly better than or comparable to that of random models, while the second cluster groups tasks leading to significantly worse neural explainability. As for voluntary movement, redundancy reduction and the tasks aimed at regressing limb kinematics including acceleration were associated with lower neural explainability. Tasks aimed at regressing the position alone were also part of this cluster, unlike in the active condition.

Since we combined data from both brain areas for our task comparison, we wondered whether the same clusters would emerge when projecting task for each brain area independently (Supp. Fig. 8B,D,F and Supp. Fig. 9B,D,F). We again observed two clusters for passive and more clusters for the active movement. Overall, the same tasks were best for S1 and CN individually, with the difference that the task of regressing posture only (HP/LP/JP) appears in the best cluster for S1 in the active case (Supp. Fig. 8B,D,F). This might suggests that S1 is less optimized for velocity coding than CN.

To summarize, we contrasted thousands of candidate models trained on 16 tasks. The most “brain-like” models were those trained to predict the kinematics of the limb from simulated spindle inputs, independently of the coordinate system. These models outperformed linear models and data-driven models.

### Deeper network layers best explain CN and S1

The models we constructed have multiple hierarchical layers. Anatomically, CN is just two synapses away from the spindles and cortical S1 area 2 another two (19, 20, 36). Therefore, area 2 may perform higher-level computations on the processed proprioceptive information such as integrating proprioceptive signals from multiple muscles or extracting more complex features. Therefore, we tested for the best models whether early model layers are more similar to CN, and deeper model layers to S1. To this end, we computed the distribution of the depth of the best selected layer among all TCN networks trained on the hand position and velocity task (Figure 6A, B). We found that deeper layers are typically the best at explaining the neural activity for both CN and S1. This was true for both active and passive trials over a wide range of network depths (Supp Fig. 10) and on a per-neuron basis (Supp. Fig. 4). As models were optimized to regress the location and velocity of the hand, deeper layers tend to develop features related to the arm kinematics which might be the reason why those layers were the best at predicting the neural activity. However, we did not find evidence that neural networks reproduce the hierarchical organization of the somatosensory system as CN and S1 are similarly explained by deep layers of the network.

**Figure. 6:**
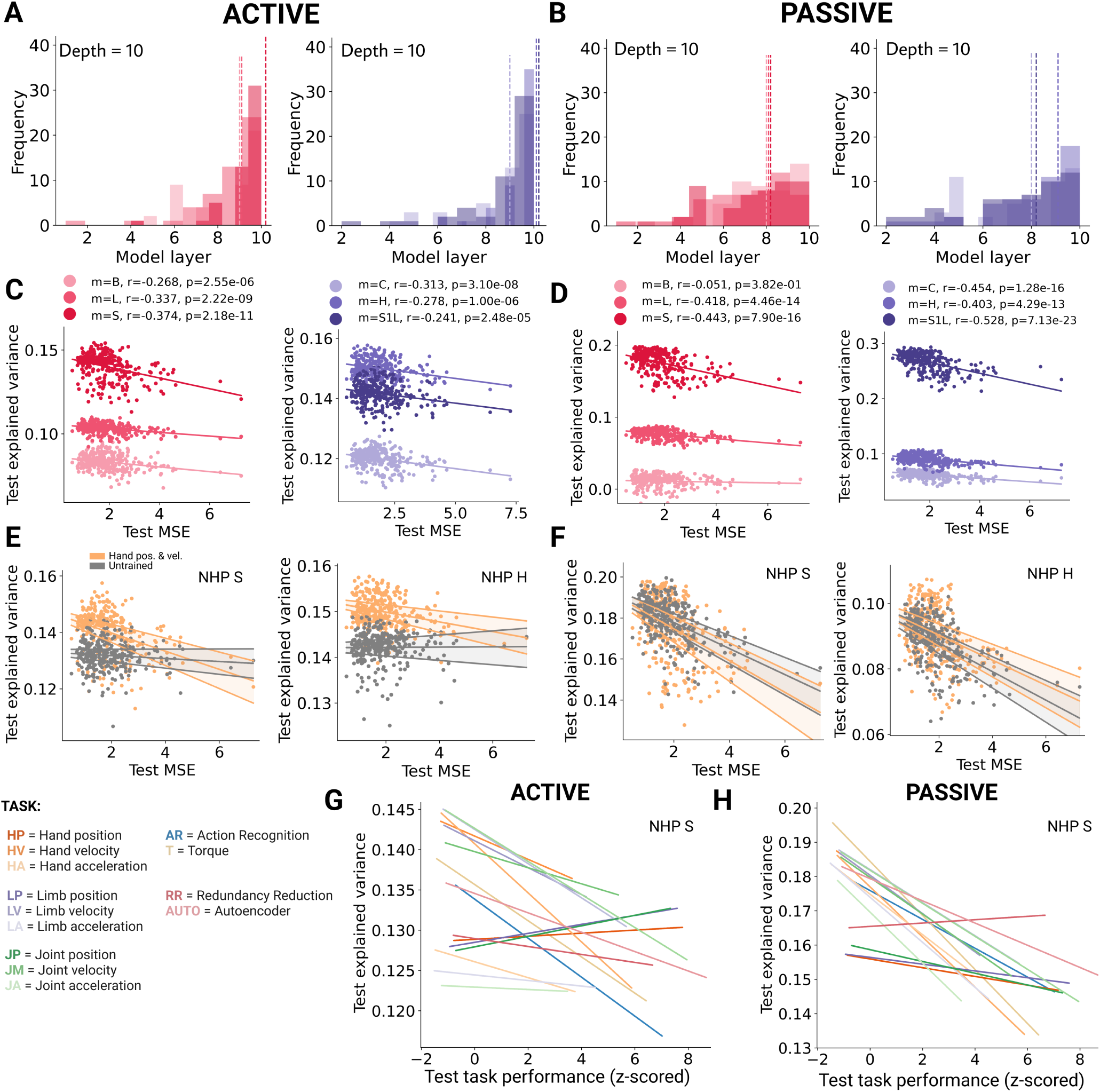
Task comparison show that kinematic-based tasks develop the most “brain-like” features and outperform untrained models. **A.** Frequency of the best-predicting DNN model layer for all models containing exactly 10 layers. The colors denote individual NHPs. The dashed lines correspond to the median best layer. **B.** Same as F for passive predictions. **C.** Scatter plot of the relationship between test explained variance on neural data, and network task performance (MSE) of each neural network in the hand position and velocity task (HP & HV). Each data point represents the neural explainability of a single DNN model. In total, N=300 DNN architectures (100 per TCN subtype) are shown. Lines indicate linear fits for each NHPs. The Pearson’s *r* correlation coefficients and the p-values are shown for each NHPs (CN m ∈ [B, L, S], left; S1 m ∈ [H, C, S1L], right), with different colors. Note that for the regression tasks, lower MSE equals better performance, hence the negative correlation coefficients. **D.** Same as C for passive movements. **E.** For example NHP S (left) and NHP H (right), scatter plots of test explained variance against model task performance for models trained on the hand localization task (orange, same data as panels C for the corresponding NHPs). In gray, scatter plots of the explained variance obtained using the untrained models against the performance on the task of their trained counterparts. (Note that the untrained models perform poorly on the task.) Lines indicate linear fits and shaded areas the 95% confidence intervals. **F.** Same as E for passive movements. **G.** Linear fits showing the relationship between the test explained variance during the active movement condition, for example NHP S, and the performance of neural networks for all tasks when evaluated on test synthetic dataset. Each color is a task as in Fig. 5A. Note that performance metrics are z-scored to compare tasks and that higher values mean poorer model performance. **H.** Same as A for passive movements.

**Figure. 7:**
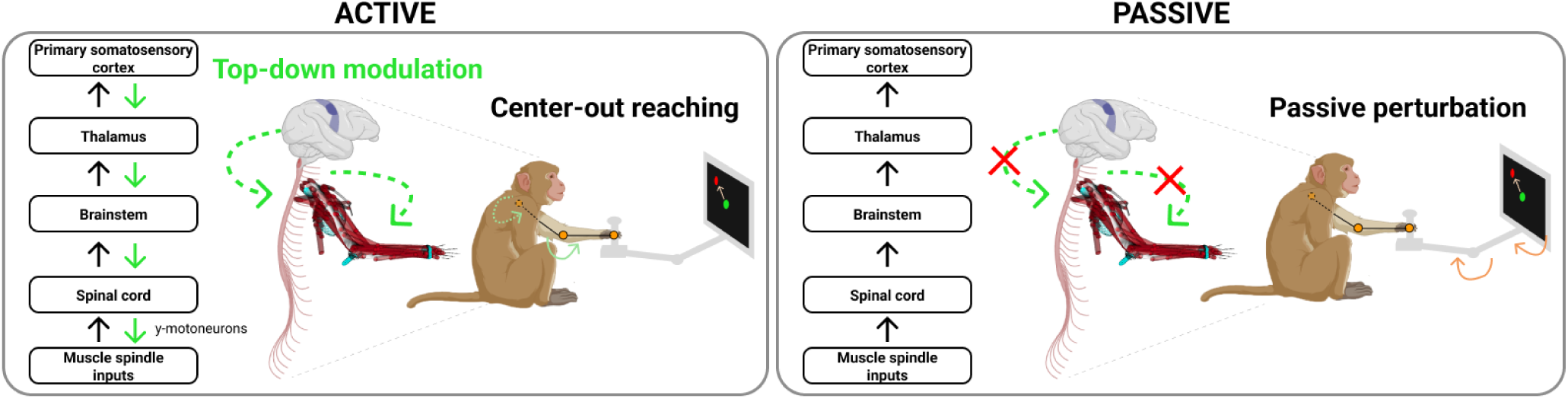
Schematic illustration of the possible top-down modulation along the proprioceptive pathway. The difference in task-driven neural predictions between active and passive movement suggest a possible top-down modulation at different levels of the proprioceptive pathway to extract relevant features for goal-directed movements.

### Task network performance strongly correlates with the ability to explain CN and S1 neural activity

For each single task, we trained a variety of neural networks that achieve diverse neural explainability (see Figure 5C,F). Different factors could potentially contribute to this variability, including the network’s architectural details, the movement condition, the task objective or the model’s task performance.

To untangle some of these factors, we first selected the most brain-like task identified in the active movement condition, e.g. the hand position and velocity task (HP & HV), and examined the relationship between task performance and neural explainability. For all NHPs, we linearly fitted the neural explainability using the task performance and found a significant correlation (Figure 6C,D). Thus, for models trained on the EgoHand hypothesis, better task performance leads to better neural explainability.

This correlation suggests that deeper models are better at explaining neural data, as they tend to be better across all tasks (Figure 2D). Moreover, we observe this correlation for passive movement data even though untrained models are not worse than trained models (Figure 6D). Therefore, we sought to isolate the effect of task-optimization by comparing the correlation between neural explainability and task performance in trained and untrained models (Figure 6E,F). To achieve this, we ordered the untrained models based on the task performance of their corresponding trained networks. For active data, explained variance consistently improved with task-optimization where task-driven models showed a higher and significant correlation than untrained models, which exhibited no correlation for nearly all NHPs (Supp. Fig. 12A incl. p-values). On the other hand, for passive data, untrained and trained models exhibited a similar correlation with task-trained performance (Supp. Fig. 12B incl. p-values). This suggests that while task-optimization is crucial for the active condition in addition to architectural parameters, the right choice of architecture alone can yield better predictions in the passive condition.

As we investigated the best “brain-like” task, we checked whether such correlation exist for the other 15 tasks. We found a significant correlation for almost all of them which means that models that are better at performing a task are also better at predicting neural activity, in both active and passive conditions (Figure 6G,H). Specifically, we observed that the most brain-like tasks had higher fitting offsets and correlation coefficients for the active condition (Supp. Fig. 11). The significant correlations for all 16 tasks, also indicates that the proprioceptive system is optimized for all those roles.

Together, our results consistently show that models that are better at performing a task are also better at predicting neural activity, highlighting the influence of both network architectural choices and task optimization. These results suggest a path for making better proprioceptive models, by identifying architectures that better solve tasks on synthetic data.

## Discussion

We expanded our normative framework combining musculoskeletal simulation and task-driven neural network modeling to compare different hypotheses about functional processing in the proprioceptive pathway (15). We trained deep neural networks, which used proprioceptive signals (muscle length and velocity) as inputs to perform 16 behavioral tasks. Importantly, we did not use any neural activity during training, but only at test time, we probed generalization. Namely, if the representations developed by task-driven models could predict the neural data from non-human primates during arm movements. Through a large-scale comparison of computational tasks and neural network models, our work shows that task-driven models outperform classical encoding models of neural activity (31, 34–36, 45) in the brain stem (cuneate nucleus) and somatosensory cortex (area 2), and also outperform the same architectures trained to predict neural activity directly.

### What is the role of proprioception?

It has been hypothesized that the proprioceptive pathway supports a range of sensory and motor-related behaviors (15, 17– 19, 37–40, 47). We defined 16 hypotheses representing some of these behaviors, and instantiated them in task objectives for training neural networks models. We contrasted the learned representations by probing the networks’ representation ability to generalize from synthetic data used for training to single-trial neural activity used for testing. Our study reveals that tasks that predict the state of the limb (position and velocity jointly) are the best at explaining the neural activity in both the CN and S1 during active movements. Similarly, the group of tasks aimed at regressing individually the position or velocity predict neural activity better than those implementing other tasks, or than untrained models.

By testing many hypotheses, our work provides normative evidence that proprioceptive neurons in CN and S1 encode the kinematic state of the limb (31, 34, 41, 48, 49). Furthermore, estimating the limb state from proprioceptive signals is also essential for feedback-based control, where the actual state is compared with the state predicted by internal models (18, 50). Interestingly, the torque-based prediction tasks does not explain CN and S1 well perhaps suggesting that the proprioceptive pathway might be more relevant for kinematic representations. However, this should be further tested with closed-loop control models.

Muscle spindles signal both muscles length and velocity. This information is processed along the proprioceptive pathway to give rise to the body schema and maps (19, 24, 25, 51– 54). Past work investigated whether sensorimotor cortex encodes body position and movement as intrinsic coordinates (e.g. joint angles and torques) (55, 56) or as extrinsic coordinates (limb position and velocity in space) (31, 57, 58) or both (41, 59, 60). We find that the choice of coordinate system does not contribute to representations that better predict neural data, as models trained to predict the state of the limb (position and velocity of only the hand, the hand plus elbow or angular position and velocity of joints) were the best at explaining the neural activity regardless of which coordinate system the regression target is represented in (egocentric vs. joint coordinates). However, we note that the behavioral workspace of the considered experiment is relatively small and differences in the coordinate system representation could emerge for a bigger workspace or more complex movements.

While limb-state estimation tasks were best to predict neural activity, other tasks yielded predictions that were no better or even worse than untrained deep hierarchical neural network models. First, in contrast with task-driven models of the ventral visual stream and auditory systems, where stimulus categorization provides the best neural representations (1, 5, 6, 9), we find that network models trained on action recognition (also a classification task) were worse than untrained models. Interestingly, this task gives rise to direction-selective units, but not position-selective unit, in the models (15).

Against our initial expectations, we also observed that the neural explainability of models trained on the torque regression task is similar to that of untrained models, which could also be a limitation of the student-teacher approach we employed for this specific task. Furthermore, training networks on multiple tasks does not always produce better representations, as in the case of the tasks that included acceleration as a regression target. Here, one explanation could be the poorer task-performance (for body localization) achieved by these network models compared to the other tasks.

Similarly, both self-supervised tasks - the autoencoder and the redundancy reduction task (BarlowTwins) - explain neural data worse than the untrained models. While autoenconding explains V1 well, it is poor for higher order visual areas (9). Zhang et al. also used SimCLR, which is most comparable to BarlowTwins, and this model could explain V1, V4 and IT as well as supervised models (9). However, BarlowTwins encourages location-invariant, but category-specific representations, which appears inconsistent with proprioception but consistent with the goal of the ventral pathway. This indicates that solely compressing the muscle spindle input statistics does not lead to brain-like neural representations. Recent work proposed a topographical autoencoder to learn the spatial arrangement of proprioceptive information in cortex (16); our work suggests that other tasks might be better suited.

### Possible top-down modulation along the proprioceptive pathway

Whether the sense of posture is mediated through central or peripheral signals is a classic debate for proprioception (61, 62). Consistently with previous analyses (31, 34, 36), the task-driven models in our study also reveal that neural predictions differ for active and passive movement. In the active case, task-driven models trained on body-state estimation can lead to more “brain-like” representations which outperform untrained models, whereas for the passive case, task-driven models do not outperform untrained models. The observation was true for both CN and S1. Indeed, models predicting the state of the body were best at explaining the neural data in both CN and S1 during active movement (Figure 5), and we found little difference between the two areas in terms of best-predicting layer (Figure 6). This was surprising as S1 is downstream of CN. How can we reconcile these results? We believe that these findings suggest that, during voluntary movement, both CN and S1 receive top-down modulation from motor areas that provide behaviourally-relevant information, such as the predicted state of the body (Figure 7). Indeed, it has been shown that S1 receives anticipatory information from the motor cortex during voluntary movements (63, 64). However, due to the unpredictable nature of the passive perturbations, this top-down modulation would have been absent. Our study suggests that CN receives predicted body state information from a forward model during goal-directed movements. This top-down modulation might alter the processing of proprioceptive information by modulating the gain and sensitivity of CN neurons for particular proprioceptive stimuli that are relevant for the behavior (49, 65). Given the multiple inputs to the CN (66), we can only speculate whether CN is directly modulated by descending cortical projections (somatosensory or motor cortex) (67), indirectly by gamma motor neurons (25, 68), by spinal presynaptic inhibition (69) or an entirely different mechanism. Nevertheless, we hypothesize that CN activity might be mainly modulated by S1, or motor cortex, as our results were highly similar for both CN and S1.

### Hierarchical processing of proprioception

In vision, convolutional neural networks trained on object recognition recapitulate the hierarchy of the ventral stream; lower, middle and higher layers best explain V1, V4 and IT, respectively (1, 5). However, less is known about the hierarchy in the proprioceptive system (19, 49). The anatomical view is that CN would be considered a “lower-order” area while S1 is a “higher-order” area in the processing of proprioceptive information. Indeed, prior studies have shown that CN neurons have muscle-like activity resembling that of their afferents, while S1 area 2 neurons possess more complex firing patterns, in part because single neurons in area 2 receive both proprioceptive and tactile inputs (49, 70), consistent with theoretical predictions related to the distribution of preferred directions along the proprioceptive pathway (15). Nonetheless, we find that deep layer representations are more accurate than early layers for both CN and S1. This was true in both active and passive movement conditions (Figure 5), although more markedly in the former, again suggesting that CN and S1 neurons may be more alike than previously thought. We do note that unlike Yamins et al. that considered three cortical areas (1, 5), we compared only two areas, only one of which was cortical. Thus, it would be an interesting comparison to see how the retina and the LGN might map onto artificial hierarchical vision models. Likewise, testing our models on thalamic signals and other proprioceptive cortical areas, such area 3a or area 5 (46, 71, 72), could help to refine how well network model layers map onto the proprioceptive pathway.

### Limitations of the study

Our models were trained using only synthetic, passively generated data. Despite this limitation, task-driven models capture the proprioceptive dynamics better than untrained models for the active conditions rather than the passive one. This is surprising as our spindle models do not include muscle force or gamma modulation. It also appears not simply to be a consequence of the amount of experimental data used for training the linear regression. However, future models will require the simulation of muscle activations and muscle force, beyond muscle kinematics alone, in order to better reflect the inputs of muscle spindles and other receptors which contribute to the sense of proprioception (19, 25). Indeed using OpenSim makes this challenging due to the slow simulation speed. We are sure that future work by us and others will leverage fast musculoskeletal models that became recently available (73). Fast models are required for learning controllers with RL (73–75) or optimal feedback control (76) for motor skills.

Furthermore, building directly on (15) we employed interpretable convolutional and recurrent network architectures to model feedforward proprioceptive processing (15). These network architectures integrate muscle inputs with spatial and temporal weight sharing, yet proprioceptive circuits are likely “non-Euclidean”. In future work, it will be important to test more general inductive biases, such as reciprocal inhibition between agonist and antagonist muscles mediated by interneurons (77) or the musculotopic spatial-map observed in the spinal cord (78). In other words, the next generation of models should better approximate the inductive bias governed by the anatomy of sensorimotor systems (79). Second, while we modeled temporal integration using temporal convolutions and LSTM cells, additional recurrent architectures could include top-down modulation (80, 81). Overall, these additional constraints might increase the explainable variance and generate additional insights into proprioception.

## Methods

### A. Behavioral and neural experimental data

#### Non-human primate center-out reaching task

We used data obtained from experiments with non-human primates (NHPs) (rhesus macaques, *macaca mulatta*) performing a center-out reaching task to quantify the predictive ability of task-driven neural network models. Detailed experimental background can be found in (36, 41, 82). Here, we briefly summarize relevant aspects. In particular, the center-out task included both passive perturbations of the arm (via a robotic manipulandum) and goal-directed active reaching. For each experimental session, the total number of rewarded active trials varied. Active movement duration varied and was on average in the range [600,900] ms.

An active center-out trial consisted in two phases: a center-hold period, in which the manipulandum is kept at a fixed, central position, and a target-reaching period, triggered by a “go” cue. In the target-reaching period, the NHP has to reach one random target direction guided by a visual cursor on a screen placed at head-level of the NHP. Approximately 50% of these trials included a 130 ms passive perturbation of the arm, which preceded the active reach, towards one random target direction using the manipulandum. Such passive movements were followed by random center-hold periods. Trials were rewarded if the NHP held the position of the cursor on the target location for a sufficiently long time after active reaches only. Target directions were in the range [0°, 270°] inter-spaced by 45°or 90°depending on the NHP behavioral session (NHP S, H, C: 8 directions; B, L: 4 directions).

#### Inferring proprioceptive stimuli from the center-out reaching task

During the center-out reaching task, 8 keypoints located from the shoulder to the hand were tracked with DeepLabCut (44, 83). These keypoints were used to generate realistic proprioceptive stimuli while passively executing reaching movements with the arm. For this purpose, we used an open-source OpenSim musculoskeletal model of the *macaca mulatta* upper limb developed by Chowdhury et al. (41) based on a previous NHP arm model by Chan & Moran (84). The model includes 39 Hill-type muscle-tendon actuators crossing the shoulder, elbow, forearm and wrist.

Using the OpenSim 3.3 simulation environment (85, 86), we scaled the musculoskeletal model to match the size of each NHP based on the distance between the markers of the Open-Sim model and the tracked markers. We used the markers trajectory to compute the trajectory of each joint by inverse kinematics. By passively moving the simulated arm model along these trajectories we computed, at each time point, the equilibrium muscle lengths **m**(*t*) ∈ R^39^ for all 39 muscles as the sum of the fiber muscle length and the tendon length. We computed equilibrium muscle configurations given joint angles as an approximation to passive movement. The muscle velocities are computed as the time derivative of the muscle lengths, using first order finite differences (time step: 0.01 s). The muscle lengths and velocities are sampled at a frequency of 100 Hz.

#### Extracellular electrophysiology recordings in proprioceptive areas

Extracellular neural activity was recorded with Utah arrays in two areas processing proprioceptive information: the cuneate nucleus (CN), located in the medulla of the brainstem, for 2 NHPs (NHPs B and S), and in Brodmann’s area 2 in the primary somatosensory cortex (S1) of 2 other NHPs (NHPs C and H), and in both of these areas in one fifth NHP with double implantation (NHP L). After spike sorting, single-unit responses (in spike counts) during center-out trials were binned in 10 ms window and firing rates were computed using a 50 ms smoothing time window (see also “Robustness to hyperparameters” section). For each NHP session data, this resulted in neural data sets of *N*_trials_ *× N*_neurons_ *× N*_time_, to be predicted by neural network model activations. In this work, we kept all but non-firing neurons (average frequency below 0.01 Hz), across all figures unless otherwise specified.

### B. A large-scale synthetic NHP proprioceptive dataset

We build on the computational framework developed by Sandbrink et al. (15) by adapting it to a musculoskeletal model of the upper limb of a non-human primate (84). Specifically, we generated a large-scale dataset of realistic proprioceptive inputs such that the range of the experimental muscle stretch lengths and velocities was a subset of the synthetic dataset. Like Sandbrink et al, we used the UCI Machine Learning Repository character trajectories dataset (87, 88) keeping only the 20 single-stroke characters but we used different spatial and temporal parameters. Since we aimed to study the proprioception of the whole NHP arm, we interpolated the trajectories to lie within a 5 × 5 cm square. In order to get the 10 ms time interval with which the firing rate was computed (100 Hz), we downsampled the trajectories from 200 Hz to 100 Hz. The resulting 2,858 character trajectories served as the basis for the end-effector (hand) trajectories. Using these end-effector trajectories, we generated realistic proprioceptive stimuli as done for the data from the center-out reaching task using the same OpenSim musculoskeletal model of the *macaca mulatta* upper limb (41). The only difference is that, while the model has seven degrees of freedom (DoF), three DoF were eliminated by enforcing the palm angle to be 0°. The four remaining DoF are the shoulder flexion (*θ_sf_*), shoulder abduction (*θ_sa_*), shoulder rotation (*θ_sr_*) and elbow flexion (*θ_ef_*). The joint angles for the four DoF are computed from the end-effector trajectories using the same constrained inverse kinematics used by Sandbrink et al. (15) but building a 2-link 4 DoF arm with arm-lengths corresponding to those of the NHP arm model (41).

While in Sandbrink et al. (15) the joint angles are defined in spherical coordinates, the joint angles of the NHP musculoskeletal models are in Cartesian coordinates. Therefore, we adapted the rotational matrices of the forward kinematics used to determine the end-effector position **e** ∈ R^3^ in an absolute frame of reference S centered at the shoulder:

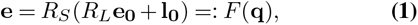

with joint-angle configuration of the arm *q* = [*θ_sf_, θ_sa_, θ_sr_, θ_ef_*]*T*, position of the end-effector **e_0_** and elbow **l_0_** when the arm is at rest, and rotation matrices

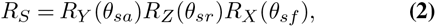

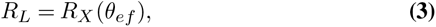

where *R_S_*is the rotation matrix at the shoulder joint, *R_L_* is the rotation matrix at the elbow joint and *R_X_*, *R_Y_* and *R_Z_* are the three rotation matrices around the *X*, *Y* and *Z* axes. The axes are defined according to the model by Chan and Moran (84), with the difference that for this model the *Y* and *Z* axes are rotated by *−*90° to match the axes defined in Opensim.

By applying the same constrained inverse kinematics used by Sandbrink et al. (15), we computed the joint-angle trajectories corresponding to each end-effector trajectory. Those trajectories were subsequently used to passively move the musculoskeletal model in OpenSim. We changed the sign of the two-link *θ_sa_* joint angle to adapt it to the corresponding OpenSim NHP upper limb model (41). Therefore, we computed, as previously explained, the muscle stretch lengths and muscle velocities at each time step for all 39 muscles considering also muscles which are not involved in the actuation of the four DoFs.

We augmented the generated joint-angle trajectories by applying the same affine transformations used by Sandbrink et al. (15) with the difference that we provided another degree of variation, i.e. changing the initial configuration of the two-link arm (see Table 1 for parameter ranges). First, we created a dataset of end-effector trajectories of 500’000 samples by generating variants of each original pen-tip trajectory, by scaling, rotating, shearing, translating and varying its speed. Second, for each end-effector trajectory, we computed the joint-angle trajectory by performing inverse kinematics. Third, we simulated the muscle length trajectories and corresponding muscle velocities. Since different characters take a different amount of time to be written, we padded the movements with static postures corresponding to the initial and final postures of the movement, and jittered the beginning of the writing to maintain ambiguity about when the writing would begin. From this dataset of trajectories, we filtered samples with muscle jerk values (third time derivative of the muscle length) higher than 1rad/s^3^ using a median filter with kernel size equal to 5 to remove OpenSim artifacts. Then, we selected a subset of trajectories such that the integral of joint-space jerk was less than 1rad/s^3^ so as to ensure that the arm movement is sufficiently smooth. Among these, we picked the trajectories for which the integral of the muscle-space jerk was minimal, while making sure that the dataset is balanced in terms of the number of examples per class, resulting in 294,711 samples. This dataset consists of muscle lengths and velocities of 39 muscles over a period of 400 time points, simulated at 100 Hz (*i.e.* 4 seconds). For each trajectory of the dataset, we saved the corresponding arm kinematics: end-effector and elbow position in egocentric or joint reference frame as well as their first and second derivatives. In this way, we could develop multiple computational tasks by training deep neural networks that take as input proprioceptive inputs (muscle length and velocity) and predict different targets depending on the task.

**Table 1:**
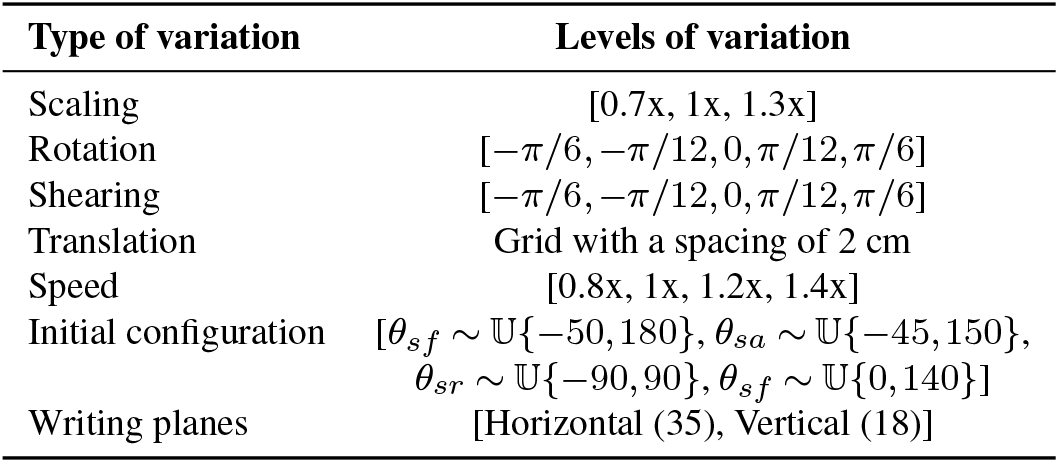
Range of the data augmentation variables applied to the original pen-tip trajectory dataset. Furthermore, the character trajectories are translated to start at various starting points throughout the arm’s workspace, the latter comprising movements in 35 horizontal and 18 vertical planes.

### C. Task-driven deep neural network models of the proprioceptive pathway

We describe here neural network models of proprioception, focusing on the following core components: network architectures, network task objectives and learning algorithm.

#### Neural network architectures

Just like Sandbrink et al. (15), we used feedforward convolutional and recurrent neural network models as architectural proxies of the proprioceptive pathway, where both architectural classes contain spatial convolutions over muscles. These deep neural networks were trained on individual tasks hypothesized to be relevant to model the sense of proprioception. The input of models comprises simulated proprioceptive inputs of muscles spindles, i.e. muscle length and muscle velocity, which are stacked together forming two channels. Specifically, the input is characterized by the following dimension [*B × T × N_muscles_ × N_channels_*], where *B* represents the batch size, *T* is the length of the trajectory which is kept fixed at 400 time points (corresponding to 4 seconds), *N_muscles_* is the number of muscles which is also a fixed quantity (39 muscles) and *N_channels_* is equals to 2 (muscle length and muscle velocity).

Each convolutional layer contains a set of convolutional filters of a given kernel size and stride, along with response normalization and point-wise non-linearity (here, rectified linear units). The convolutional filters can either be onedimensional, processing only spatial (aka muscle) or temporal information, or two-dimensional, processing both types of information simultaneously. We sampled 100 TCNs for each type (spatial-temporal, temporal-spatial and spatiotemporal) as well as 50 Spatial-LSTMs (Sample space hyperparameters see Table 2 and 3)). More details can be found in Sandbrink et al. (15).

**Table 2:**
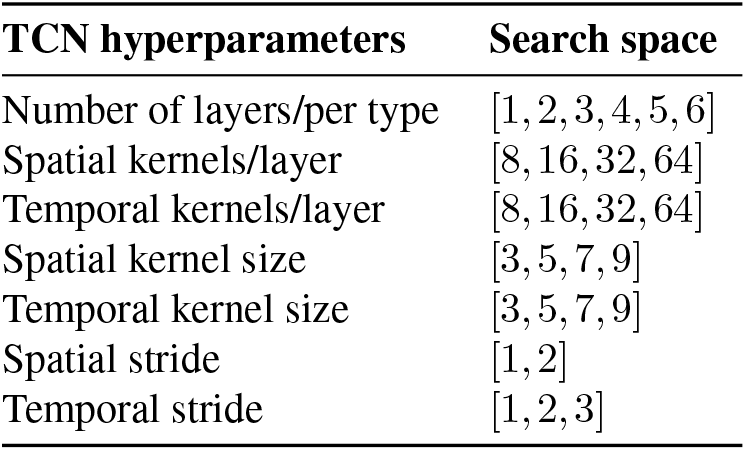
Hyperparameters to search TCNs. First, a number of layers (per type) is chosen, in the range 2-8 (in multiples of 2) for spatial-temporal and temporal-spatial models and 1-4 for the spatiotemporal ones. Next, a spatial and temporal kernel size is picked (per layer), which remains fixed throughout the network. For the spatiotemporal model, the kernel size is equal in both the spatial and temporal directions in each layer. Then, for each layer, a number of kernels (or feature maps) is chosen such that it always increases along the hierarchy. Last, a spatial and temporal stride are chosen. For each network subtype, 50 models are randomly sampled using these hyperparameters. The same sampling strategy is applied for model with 10/12 layers with the difference that the temporal and spatial stride is in-homogeneous among the layers. Another 50 models are randomly sampled, for a total of 100 models per architecture subtype. Table adapted from (15).

| TCN hyperparameters | Search space |
| --- | --- |
| Number of layers/per type | [1, 2, 3, 4, 5, 6] |
| Spatial kernels/layer | [8, 16, 32, 64] |
| Temporal kernels/layer | [8, 16, 32, 64] |
| Spatial kernel size | [3, 5, 7, 9] |
| Temporal kernel size | [3, 5, 7, 9] |
| Spatial stride | [1, 2] |
| Temporal stride | [1, 2, 3] |

#### Neural network task objectives

A central hypothesis in goal-driven modeling is that sufficiently rich and ethologically relevant tasks are required to constrain a model of the sensory pathway that is thought to support these behaviors. Using the same network architectures, we train deep neural networks on several supervised and unsupervised objectives, which are summarized in Table 4. For all tasks the inputs are identical and are given by the muscle length and velocity trajectories.

**Table 3:**
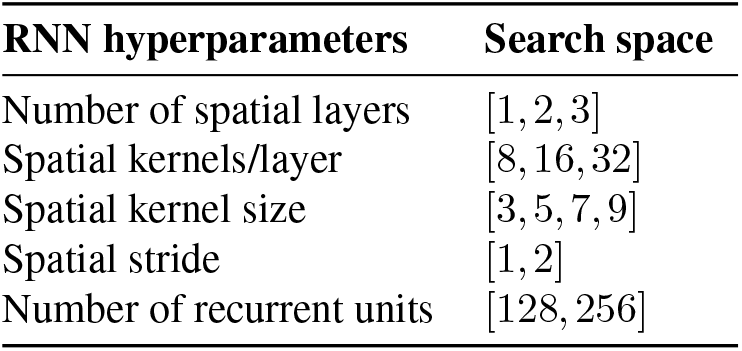
Hyperparameters to search RNNs. A number of spatial convolutional layers is first selected then a number of recurrent units.

| RNN hyperparameters | Search space |
| --- | --- |
| Number of spatial layers | [1, 2, 3] |
| Spatial kernels/layer | [8, 16, 32] |
| Spatial kernel size | [3, 5, 7, 9] |
| Spatial stride | [1, 2] |
| Number of recurrent units | [128, 256] |

**Table 4:** List of neural network task objectives divided in supervised tasks and unsupervised tasks (pos: position; vel: velocity; acc: acceleration).

| Acronym | Task description |
| --- | --- |
| Supervised tasks |  |
| HP | Hand position<br>(End-effector pos.) |
| HV | Hand velocity<br>(End-effector vel.) |
| HP & HV | Hand position and velocity<br>(End-effector pos. and vel.) |
| HP & HV & HA | Hand position, movement and acc.<br>(End-effector pos., vel. and acc.) |
| LP | Limb position<br>(End-effector/elbow pos.) |
| LV | Limb velocity<br>(End-effector/elbow vel.) |
| LP & LV | Limb position and velocity<br>(End-effector/elbow pos. and vel.) |
| LP & LV & LA | Limb position, velocity and acc.<br>(End-effector/elbow pos., vel. and acc.) |
| JP | Joint position<br>(Joint angles) |
| JV | Joint velocity<br>(Joint angular vel.) |
| JP & JV | Joint position and velocity<br>(Joint angles and angular vel.) |
| JP & JV & JA | Joint position, velocity and acc.<br>(Joint angles, angular vel. and acc.) |
| AR | Action recognition<br>(Character recognition) |
| T | Torque<br>(Joint torques) |
| Unsupervised tasks |  |
| RR | Redundancy reduction<br>(Barlow twins) |
| AUTO | Autoencoder |

#### Redundancy reduction (Barlow Twins)

The redundancy reduction task, also named Barlow Twins, is a self-supervised task. The neural network is trained to develop invariant representations given two distorted versions of the same input as proposed in computer vision (43). In the sensorimotor case, we generated two distorted inputs from a muscle input trajectory by randomly masking two temporal windows of 500 ms (one at the first half and the other at the second half for a total of 1000 ms over 4000 ms of the entire trajectory). These two distorted inputs are fed into the network generating two different representations (*Z_A_*and *Z_B_*) and the network is trained to make the cross-correlation matrix between the two representations match the identity matrix. In this way, the latent representations of two distorted inputs are forced to be similar, while minimizing the redundancy between them. A projector of 3 fully connected layer of 256 units each was added to the previously described network to match the implementation described in the original work (43).

The networks minimize the Barlow Twins loss (43):

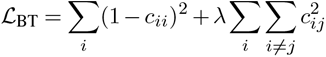

where *c* is the cross-correlation matrix between the two representations defined as:

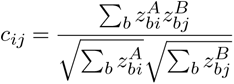

and where *λ* = 0.005, *b* represents the batch sample and *i*, *j* represent the vector dimension of the representation.

We assessed the quality of the representations of the networks trained on redundancy reduction in terms of the model’s ability to transfer to other tasks (which are easier to interpret). To compute task-transfer performance, the weights of the network are frozen and a linear readout is trained to solve the action recognition task or the hand-localization task (using the same training data). We found networks can achieve 50 % classification accuracy and around 7 cm errors for the localization of the hand (not shown).

#### Autoencoder

In order to reconstruct the input as unsupervised task, we added a symmetric decoder comprised of transposed convolutions and a linear readout. The autoen-coder networks were trained to minimize the mean square error between the original and reconstructed input. For interpretability, performance is also illustrated as relative error (Figure 2E) and broken down in muscle length as well as velocity (Supp. Fig. 2D).

#### Action recognition task

Given that the synthetic dataset comprises passive movements corresponding to different Latin characters in the 3D space, we trained the networks to classify the character. The networks are trained by minimizing the softmax cross entropy loss. Therefore, the network is trained to recognize the same actions performed in different locations.

This task was proposed in Sandbrink et al., and was found to give rise to uniformly distributed direction selective units along the proprioceptive hierarchy (15). Similarly, we also considered the trajectory decoding task, which we generalized to include velocity and acceleration as well as considered different coordinate frameworks.

#### Regression tasks (State estimation in different coordinate frameworks)

For each task, the network was trained to regress the target quantity. In particular, we developed twelve regression tasks. For the “EgoHand”, “EgoLimb” and “JointLimb” hypotheses we developed the following behavioral tasks: (1) regressing the position alone, (2) regressing the velocity alone, (3) regressing both position and velocity, (4) regressing simultaneously position, velocity and acceleration. Specifically, the networks were trained to regress the end-effector/hand for the “EgoHand” hypothesis, the end-effector and the elbow for “EgoLimb” hypothesis. Here, the “Ego” prefix denotes that the target is represented in egocentric coordinates, whereas the four limb joint angles/velocities/accelerations were regressed for the “JointLimb” hypothesis. All networks for the regression tasks were trained to minimize the mean square error (MSE). Neural networks can have different temporal strides which means there is a temporal dilation and the overall time range of model activations could be different from the one of the input. Therefore, we downsampled the target trajectory to match the corresponding time resolution of the model activations of the final layer.

#### Joint torque regression

OpenSim is relatively slow for efficiently learning to predict controllers to reach arbitrary goal locations (73). Thus, we designed a student-teacher approach. More specifically, we trained a reinforcement learning agent to perform a center-out task in the 3D space, using the efficient physics simulator PyBullet (89). A policy network was trained with the Soft Actor-Critic (90) algorithm to control a robotic arm scaled to a similar size as the primate arm. The model learns to predict joint torques provided the current proprioceptive state as well as a target with a frequency of 100 Hz. However, PyBullet does not support the simulation of musculoskeletal models, meaning that the proprioceptive state of the arm only included joint angles and velocities. This is in contrast with the other task-driven networks presented in this work, which receive the muscle state as inputs. Therefore, we used the trained policy network as a *teacher*, to train the *student* networks (TCNs and Spatial-LSTMs) via knowledge distillation (Supp. Fig. 1). We used the trained agent to generate a dataset of 100.000 center-out reaching trajectories, including joint angles, joint velocities and joint torques. The dataset was built sampling randomly targets in the whole working space of the arm. As described in the previous sections, we used these joint angles to simulate muscle length and velocity by computing equilibrium muscle configurations. In this way, the input data represents the proprioceptive muscle state coherent with the other tasks. Therefore, we created a large dataset to train different neural networks to control the arm and perform reaching movements, using the same input type as well as the same encoder architectures trained for the other tasks. Since the encoder only takes muscle length and velocity as an input, we needed to also input a goal location. We provided the student network with the target by concatenating the target vector with the latent proprioceptive representation at the end of the encoder (Supp. Fig. 1A). This latent representation goes through a decoder comprising two fully connected layers to regress the joint torques by minimizing the MSE.

#### Learning algorithm, training protocol and evaluation

Neural network models were optimized using backpropagation. We used the Adam optimizer with an initial learning rate of 0.0005, batch size of 512 (0.001 learning rate and 70 epochs for the LSTM), except for the redundancy reduction task for which we used a batch size of 256, a learning rate of 0.005 for 25 epochs (0.001 learning rate and 40 epochs for LSTM). When the validation loss has not improved for 5 consecutive epoch, we retrieve the checkpoint corresponding to the best validation loss and decrease the learning rate by a factor of ten. After the second time this occurs, we end the training and the accuracy of the networks is evaluated on the test set. Importantly, each architecture was trained on all 16 tasks from the same randomly initialized state. We refer to those models as *untrained* models.

### D. Task-driven neural network models of proprioceptive neurons

#### Using center-out reaching NHP data as test stimulus

We created for each experimental session, a dataset of estimated proprioceptive stimuli during the reaching movements. We aligned the movement onset for each trial and zero-padded the input to avoid boundary effects. In this way, we used these experimental proprioceptive stimuli as test input to the frozen neural network and computed the corresponding time-varying activations.

Principal component analysis was performed on the layer activation sets to reduce the dimensionality to 75 principal components and to keep the number of features fixed across layers. We note that this is similar to the number of inputs for the muscle lengths and velocities (i.e., 2 *×* 39). We showed that the corresponding neural explainability is robust to this number (see also “Robustness to hyperparameters” section, and Supp .Fig. 4A). These components are referred to as “activations”, and served as the basis to model single-neuron firing rates recorded during the center-out reaching experiments.

#### Single-neuron predictivity models

In the following, we describe how we used neural network activations generated by NHP proprioceptive inputs to model single-neuron firing rates along the proprioceptive pathway.

We model the activity of biological neurons performing a linear regression from the PCs of each layer activation. The main reason is that it is unclear whether an exact matching of single neurons from the same area (5) exists, and there is a degree of variability in the properties of single neurons and inter-area connectivity within the proprioceptive pathway across different animals. To this end, we linearly combined model features (PCs) to form a feature set that can be used to explain single-neuron activity in CN and S1 neurons.

For each session dataset, we focused only on the time bins corresponding to actual reaching movements of the NHP (from the movement onset to the trial end). We divided the behavioral data for each NHP in 80% train and 20% test set based on the trial index. Importantly, we used the same split for all analysis, i.e. when comparing linear encoding models, task-driven and data-driven models.

For each set, the single-neuron firing rate counts corresponding to these bins were concatenated across trials (N° neurons *×* the concatenated test trials duration). Then, we used PCs of layer activations to regress the firing rate of each neuron through ridge regression. Since we used neural networks with different temporal strides, the model activations might have a temporal dilation and an overall time range which is different to the one of the neural firing rate. Therefore, we linearly interpolated the PCs of the temporal-dilated model activations to match the corresponding time range and resolution of the neural firing rate.

Proprioceptive information is continuously provided to CN and S1 during movement, but we did not fit a new linear model independently for each time bin. Instead, the objective is to learn only one linear model, i.e. the weights remains fixed across time. This approach was similarly used in Nayebi et al. (10). Therefore, we seek to find unique weights *w_n_*for each artificial principal component feature *n* at time *t* of network layer 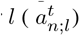 during the motion. The predicted response for neuron i at time *t* is thus:

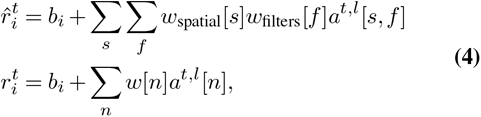

where *b* is an estimated baseline firing rate, *w* are the temporally-fixed parameters [*w*_spatial_, *w*_filters_].

For each neural network model layer, linear regression models were fitted independently for each single neuron in CN and S1. Regression weights for activations were fitted on the concatenated train session trials, and evaluated on the concatenated test session trials. The ridge regression regularization hyperparameter was validated using 5-fold crossvalidation (trial-based) on the training dataset. A final model was fitted using the optimal regularization hyperparameter on the whole training dataset.

Similarly, we used the cross-validated ridge regression model to establish a baseline encoding model of neural activity. To this end, we fitted single neural activity using kinematic variables as regressors (See also “Baseline linear encoding models of neural activity” section).

#### Neural explainability metrics

The neural explainability was quantified using the fraction of explained variance (EV) of single-neuron linear model predictions in neural responses:

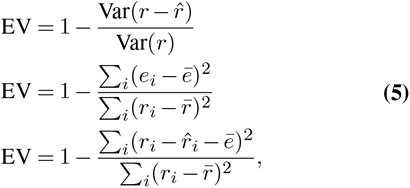

where *r* and *r̂* are the observed and predicted single neuron vector responses, *r_i_* and *r̄* are the individual observations and the mean firing rate. We note that the explained variance of a linear regression model equals the coefficient of determination, *R*^2^, when the mean of residuals *ē* is null. Thus, for each neuron, we obtained one EV score per model, per model layer and per task. To compare the neural explainability across different models, we defined the neural *explainability* which gives a sense of how good one model is at predicting the neural activity. Specifically, it is defined as:

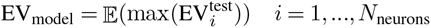

which is the average maximum EV (from the best predicting layer on the training set) for each neuron. Using this metric, we make no commitment as to which network layer should better match which brain area. Rather, we consider this way that each trained model contains layers which constitute abstract candidate locations for the most brain-like stage during proprioceptive processing, and the stage may differ between neurons of the same area.

#### Estimating neuron trial-by-trial variability

To compare single neurons and neuron populations, we quantified the trial-by-trial variability in single neuron firing rates and computed an estimate of the neuronal reliability (Supp. Fig. 3E,F). This is required since neurons are inherently noisy and the recording quality might vary across different populations. Similar to the method in (8), the estimate corresponds to the Pearson correlation coefficient between single trial firing rate and the trial-averaged firing rate for a given movement direction, averaged over movement direction conditions. Single-trial firing rates were summed using 50 ms windows, like for task-driven neural predictions. Since trials have different length during active reaches, we compute single-trial correlations using spike counts from movement onset to trial end.

#### Baseline linear encoding models

Using the same train/test trial splits of the ridge regression, we computed neural predictions using as regressors the different behavioural variables, i.e. muscle kinematics, joint kinematics and end-effector kinematics, in both Cartesian coordinates and the vector norm. We used these variables alone or in combination, to predict the concatenated trial activity of each CN and S1 neuron. Furthermore, we performed the same neural predictions with generalized linear models with Poisson distribution (Figure 3F and Supp. Fig. 3D). The performance of these baseline models was also quantified using the explained variance on test trials for each neuron. To compare with a model’s neural explainability, we computed the mean explained variance across neurons.

#### Relationship between task-performance and neural explainability

To better understand the impact of task-optimization, we studied the relationship between task-performance and neural explainability (Fig. 6). We performed the same analysis but comparing the correlation across different tasks for the same NHP (Supp. Fig. 11). Since each task has a different metric to evaluate the performance of the network, we z-scored each task-performance to make them comparable. Specifically, we took the reciprocal of the classification accuracy for the action recognition task to ensure that slopes have all the same orientation. Moreover, we also compared the same correlation both for the neural explainability obtained with task-trained networks and the one with untrained models. For the latter, we kept the task-performance of the corresponding trained models but replaced the task-driven prediction with the untrained one (Supp. Fig. 12). We assessed the robustness of the difference between the two correlation by computing the 95% confidence interval via bootstrap (N=1000) for both active and passive condition for each NHP. We observed a significant difference in correlation between task-trained and untrained models for the active case but not for the passive one underlying again the importance of task-optimization to explain neural activity for goal-directed movements.

#### Constructing low-dimensional embeddings of tasks via UMAP

We wanted to assess the impact of task-optimization both across NHPs and brain areas, while considering various neural network architectures. To achieve this, we utilized a low-dimensional UMAP (91) space to visualize and analyze the difference in neural explainability between task-driven models to their untrained counterparts (Figure 5B,E, Supp. Fig. 8 and Supp. Fig. 9)). Specifically, we computed the difference in neural explainability between a task-trained model and the corresponding untrained model. We have 300 TCN models per task, except 295 models for the autoencoder task. Therefore, we kept the maximum number of models that were shared across all tasks and NHPs (N = 292), which can be evenly divide into four groups. For each task, we randomly subsampled four groups of networks. In this way, the features of one datapoint (per task) are represented by the task-trained vs. untrained difference of the group of networks, which is thus less dependent on particular architecture parameters of a single model (N = 73). Each model is used to predict the neural activity of each NHP (N = 6). Therefore, we have 24 datapoints (4 groups x 6 NHPs) per task when comparing across NHPs, whereas we have 12 datapoints (4 groups x 3 NHPs) per brain area when comparing CN and S1 independently. This data was visualized in 2D with UMAP. We additionally validated that similar clusters emerged using PCA (Supp. Fig. 8 and Supp. Fig. 9).

### E. Data-driven models of proprioceptive neurons

We wanted to understand whether the better neural predictability achieved with task-driven models is only a consequence of the higher computational capacity of deep neural network. Therefore, we compared the neural predictions obtained with task-driven networks with the neural predictions that can be obtained by training the same neural networks to directly regress neuron firing rates from NHP proprioceptive stimuli. We refer to these models as “data-driven” models, as opposed to “task-driven” models.

Importantly, we trained these neural networks using the same training/test split of the ridge regression. In this case, we give as input to the deep neural networks the estimated proprioceptive stimuli and we train it to regress single-neuron firing rates. Overall, we have the same *N* = 300 TCN networks (used for task-driven modeling) trained on each NHP.

#### Data-driven networks training and evaluation

Each neural network model was initialized using the same initialization of the corresponding task-driven models and trained end-to-end to predict single-trial firing rates from proprioceptive stimuli. This was done per CN/S1 neuron population: the models were optimized to predict the neural dynamics of all neurons jointly, where each node of the output layer corresponds to one neuron. Consequently, the convolutional layers of the encoder are shared between neurons, thus modeling the pathway up to a particular brain area. The networks were trained to minimize the variance of the residual, i.e. the mean square error between the predicted firing rate and the ground truth. Since the movement reaching is only a subset of the entire temporal dimension of the datapoint (T = 4 s), we masked the loss in time to take into account the firing rate of the neurons related only to the movement reaching part. Overall, the loss was taken over the batch trials and averaged over neurons:

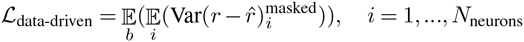

where *r* and *r*^ are the observed and predicted single neuron vector responses, and *b* represents the batch sample. Training performance of the networks were monitored by computing the mean explained variance only on the movement reaching part of the trial input:

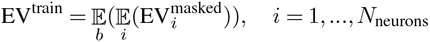

where *b* represents the batch sample. After convergence of the loss function, data-driven models are evaluated on test trials, computing the explained variance of the model neural predictions over the trial-concatenated bins corresponding to actual movement reaching data, from all test trials and for each neuron separately. As a result, neural predictions between data-driven and task-driven models are directly comparable.

### F. Robustness analysis

#### Robustness to bin width and latency

To predict single unit dynamics (Figure 4), we used a 50 ms bin width and 0 ms latency between neural data and movement (see Methods). However, we determined that task-driven models outperform linear models for across various bin widths (Supp. Fig. 13) and a wide range of latencies around 0 (Supp. Fig. 14). Therefore, we maintained these parameters throughout the manuscript.

#### Robustness to hyperparameters

The number of PCs was chosen to avoid over-fitting in the neural predictions, as it is lower than the number of units in the layer (Supp. Fig. 4A). The reasons to use principal components are two. First, the number of artificial units changes depending on the depth of the layer and the network architecture. By mapping activations to a fixed number of principal components, we remove possible biases given by the different size of the features space. Second, we do not assume a strict, one-to-one mapping of model unit (artificial neuron) to biological neurons. Rather, we hypothesize that these principal components constitute a feature space of activity that constitutes a linear basis of a neuron’s neural activity. The same procedure was also employed to compute neural predictions from randomly initialized, that is, untrained neural network models (Supp. Fig. 4B), which can then be compared to the task-trained neural predictions.

We also highlight the fact that for some NHPs, neurons might not be entirely proprioceptive but they can also carry tactile information or lower limb information (neurons in gracile nucleus, as for NHP B). Last, differences across NHPs may also reflect intrinsic variability in experimental datasets, neuronal properties, signal-to-noise ratio and total number of session trials.

We also cross-validated the neural predictions obtained when varying the size of the time window used to calculate the spike firing rates. For three models trained on the hand position and velocity task, we computed neural predictions using windows in the range [10, 100] ms, in steps of 10 ms. We settled on an intermediate window of 50 ms (Supp. Fig. 13). Similarly, we cross-validated neural predictions when shifting the time of the neural activity relative to the model activations generated from muscle spindle inputs, with latency shifts in the range [−400, 400]ms (Supp. Fig. 14).

#### Robustness to length and number of trials

Task-driven models can better explain the neural activity in the active conditions rather than the passive one. The main differences between the active and passive condition are the number of trials (Fig. 3) and the trial length (variable for active and 130 ms for passive). Therefore, we assessed the robustness of the results between the two conditions by performing the neural predictions for the active condition but decreasing the number of trials to match the number of passive trials. Moreover, we randomly selected sub-interval of 130 ms (matching the length of passive trials) because we wanted to make sure the difference was not specific to any part of the movement but could be related to any part from the movement onset to the end of the trials (Supp. Fig. 6D,E). We found that the difference between trained and untrained models is maintained for the subsampled data, thereby discarding the amount of the data as possible cofactor.

## Acknowledgments

We thank Mackenzie Mathis, John Kalaska, Sliman Bensmaia and members of the Mathis Group for helpful feedback as well as to Daniel Cavaleri for contributing to the design of the robotic arm PyBullet model used for the motor control task. We are grateful to Lucas Stoffl, Adriana Rotondo for comments on an earlier version of the manuscript.

## Funding

This work was funded by Swiss SNF grant (310030_212516) and EPFL. A.M.V.: Swiss Government Excellence Scholarship.

## Declaration of interests

The authors declare no competing interests.

---

**Supplementary Figure 1:**
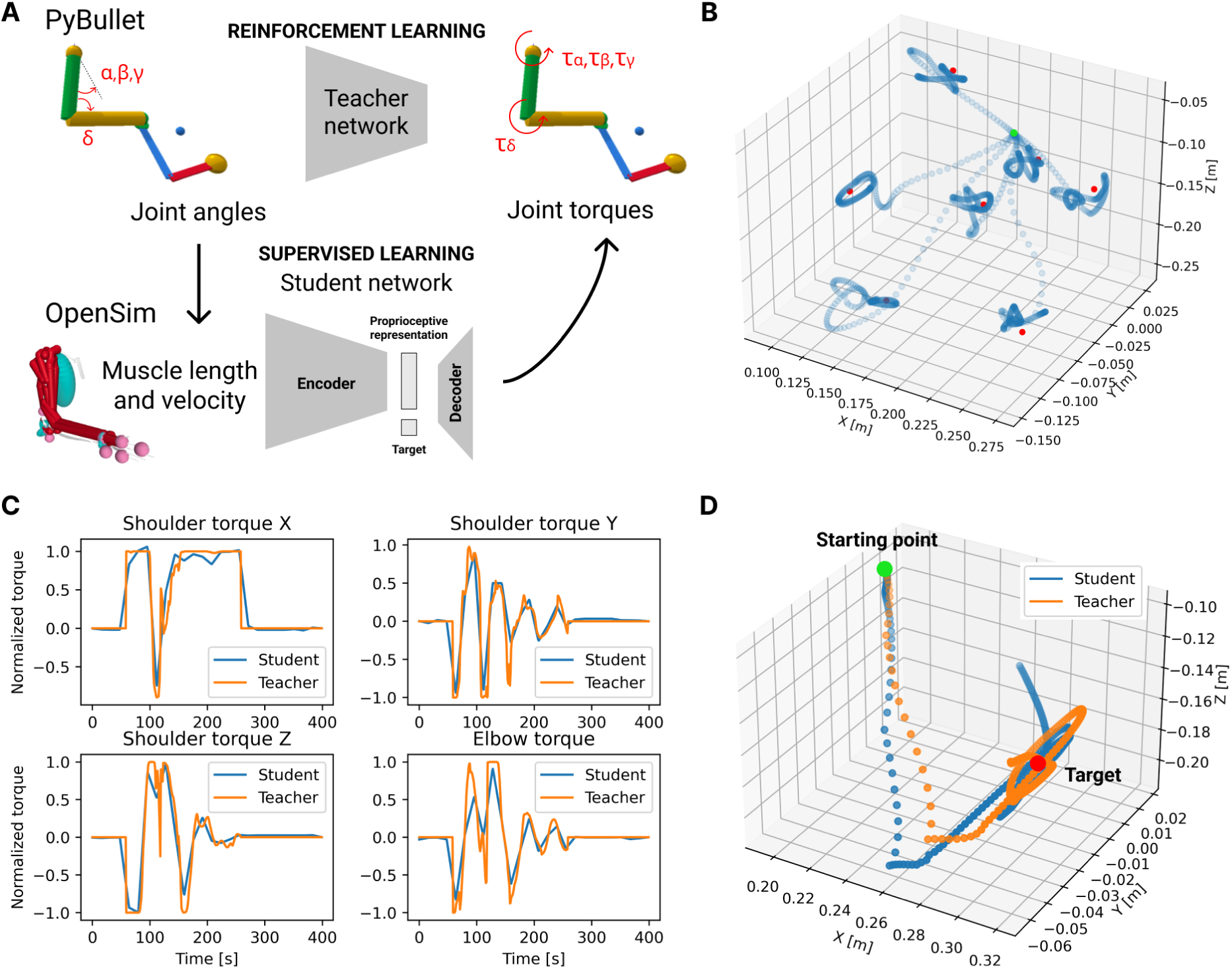
**A.** Schematics of the teacher-student training procedure to distill the task-driven neural network from a control policy (see methods). **B.** Example trajectories of hand position (blue) when the trained agent tries to reach different targets (red dots), starting from the same initial position (green dot). **C.** Comparison between each component of the torque, output by the teacher and by the student for one test episode. Despite the lower output frequency of the student network, the curves show an evident similarity. **D.** Example trajectory of the hand position, when the arm is controlled by the teacher network (orange) or by the student network (blue). Also the student network succeeds in reaching the target position (red), starting from the same initial position (green) of the teacher.

**Supplementary Figure 2:**
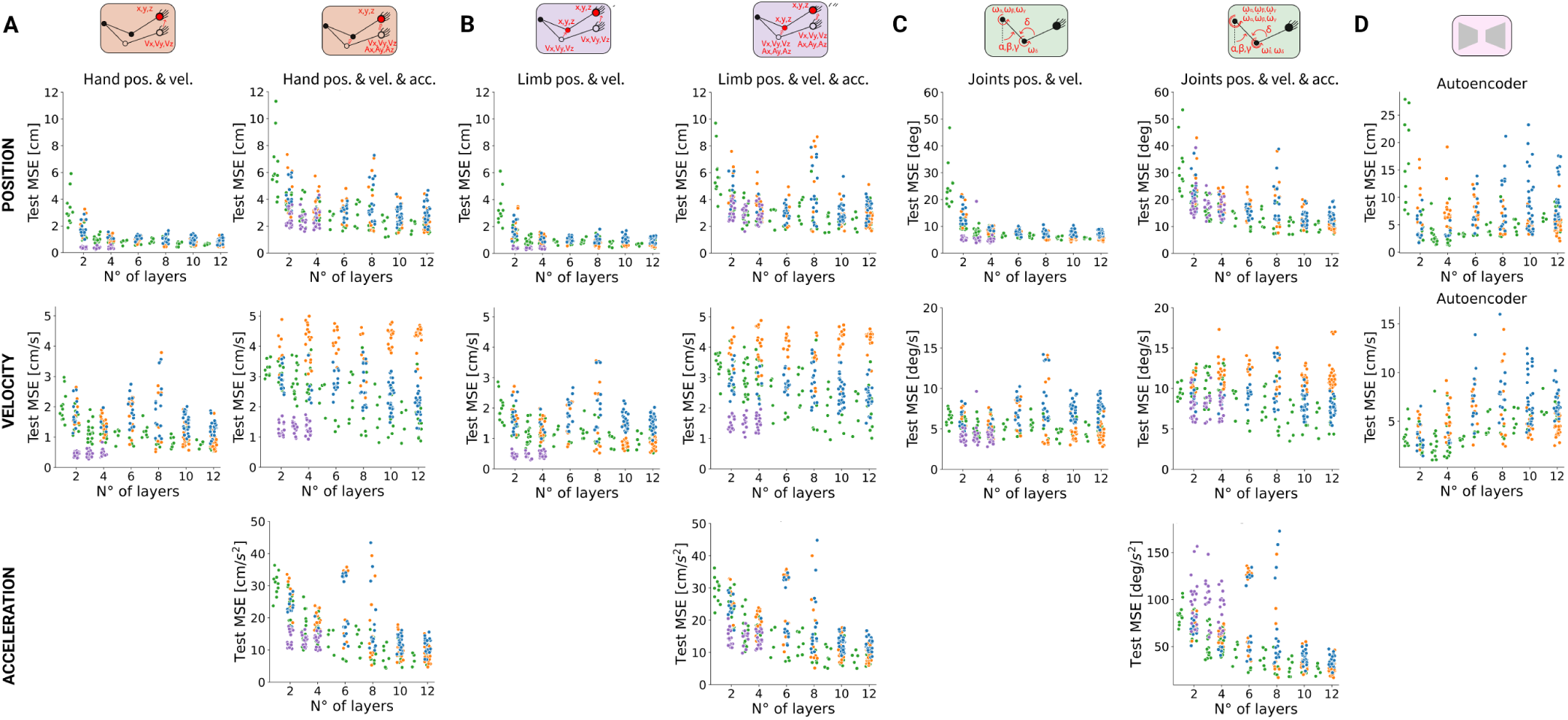
Test performance of each network model with respect to the number of layers (i.e. model depth) on tasks that combine position and velocity or position, velocity and acceleration (N=350 models per task). For each task, the test performance is decomposed in position (first row), velocity (second row) and acceleration (third row, if present). Each columns represents the combined tasks for **A.** the EgoHand hypothesis, **B.** the EgoLimb hypothesis, **C.** the JointLimb hypothesis and **D.** autoencoder networks. Performance is quantified with the mean square error (MSE). Colors indicate neural network architecture type: spatiotemporal (green), spatia-temporal (blue), temporal-spatial (orange), spatial-LSTM (purple). Task that included acceleration in the target show lower performance on the single target alone (e.g. position or velocity alone) compared to networks optimized to predict the single target.

**Supplementary Figure 3:**
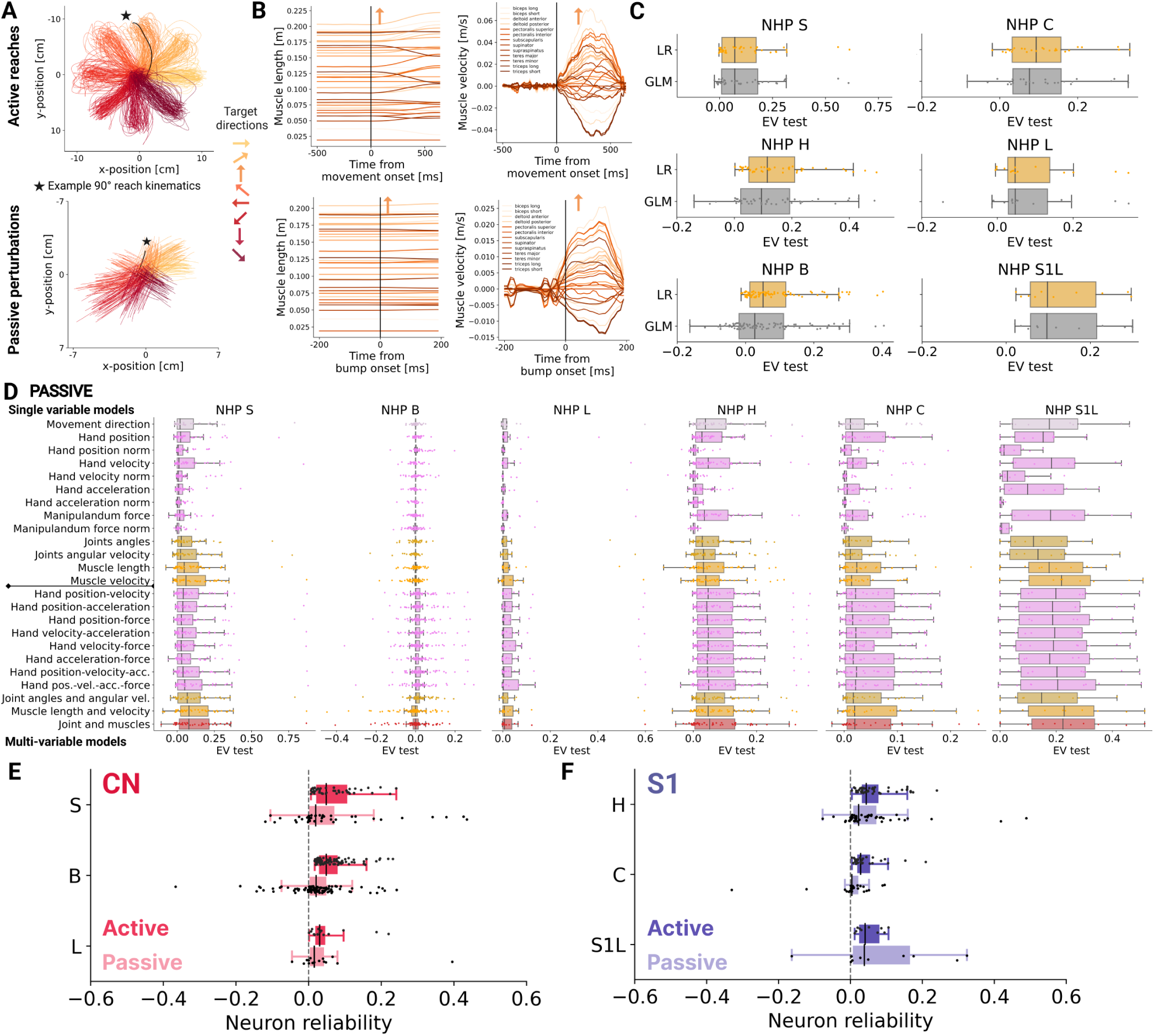
**A.** Example NHP hand trajectories during the center-out reaching task during active movements (reaches; left) and passive movements (bump perturbations; right). Each line corresponds to the center-out trajectory of one trial. The color code corresponds to movement directions. **B.** Example muscle kinematics for a single trial (indicated in panel A with a black star at the trajectory end) in both active reaches and passive perturbations, aligned at movement/bump onset. Top: muscle stretch length for selected examples muscles. Bottom: muscle stretch velocity for the same example muscles. **C.** Distribution of test explained variance between ridge regression (orange) and generalized linear model with Poisson process (gray) for each NHP using muscle length and velocity as regressors for active movements. Ridge regression showed more robust results compared to GLM, therefore we used the former in our analysis referring to those as “linear models”. **D.** Raw performance of linear tuning models fitted on task-related single-variable and multi-variables for each NHP (NHP S, L, B recorded in CN; NHP H, L, C recorded in S1) during passive movements. Individual dots represent single neurons. Task variables are color coded based on movement direction, hand, joint, and muscle kinematics. **E.** Distributions of single-neuron reliability (or self-consistent explained variance), for active and passive movements, for each non human primates in CN. Reliability is computed as the average variance of single trial firing rates explained by the movement direction specific trial-averaged firing rates. This average is calculated for each movement direction (4 or 8) and we report the average over movement directions. **F.** Same as in D but for S1 NHPs.

**Supplementary Figure 4:**
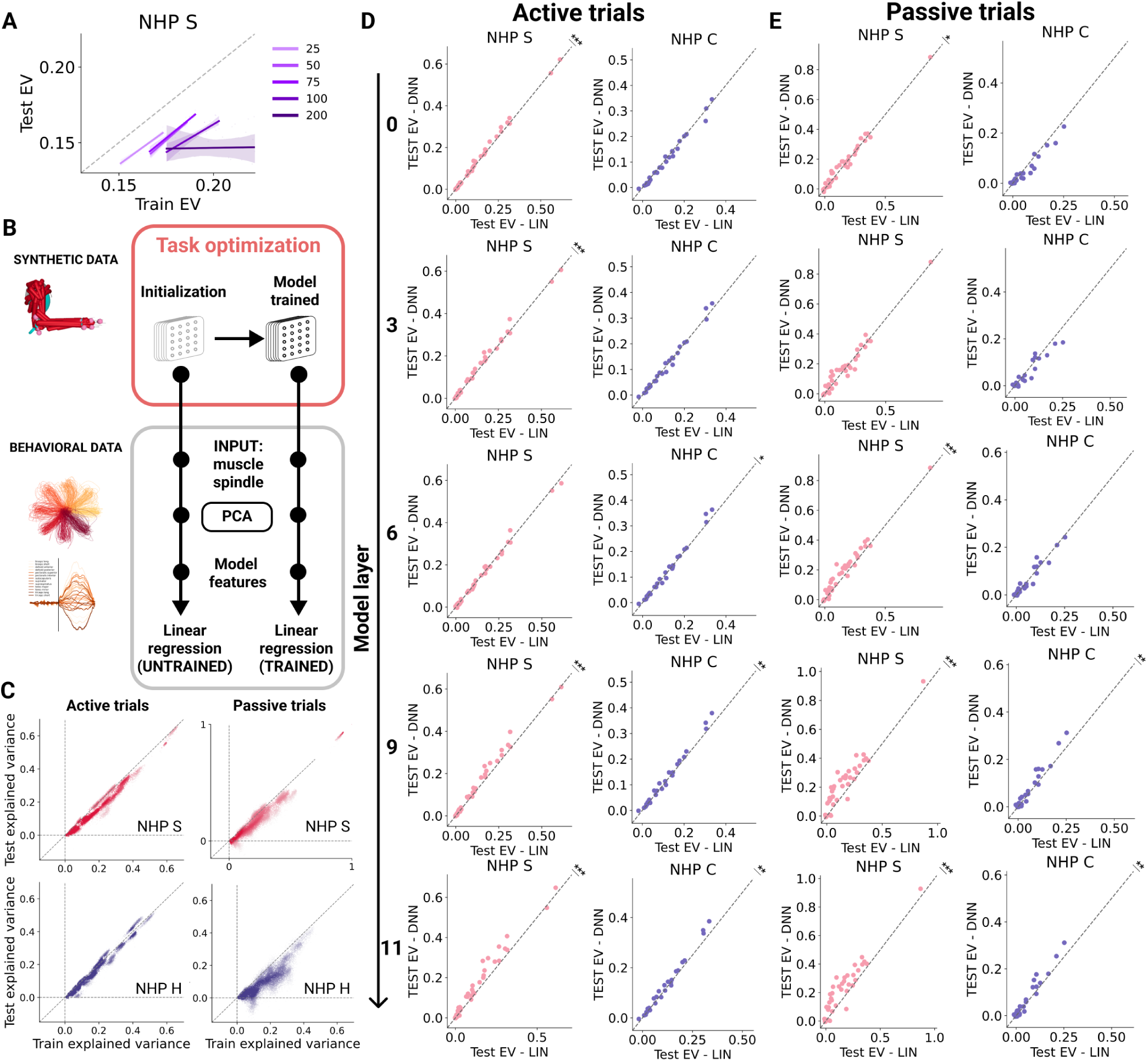
**A.** Correlation between test and train neural explainability (NHP S; CN) with respect to the number of PCs for representative models trained on the action recognition task (N=6 models, two per TCN subtype). Shaded are is the confidence interval at 95%. We note that we used 75 principal components throughout the manuscript (for fair comparison with the number of proprioceptive inputs) as it avoids overfitting but higher test EV is possible with around 100 dimensions. **B.** Neural predictions from TCNs workflow. Top: task optimization of neural network models is done using the large-scale synthetic proprioceptive input data (derived from pose estimation and musculoskeletal modeling). Bottom: experimental proprioceptive inputs during center-out reaching of NHPs is used as new inputs to the frozen neural network models to generate model activation per layer, which are combined using principal component analysis. This constitutes model features, which we generate for both untrained, random models and task-trained models. From these features, we generate neural predictions using spatial linear regression (i.e., weights do not change for each time bin). **C.** Distributions of explained variance scores between training and testing trial datasets for each NHP S (CN, top row) and NHP H (S1, bottom row) obtained from models trained on the hand position and velocity task for the active condition (left column) and passive one (right column). For visualization, N = 20000 randomly sampled single-neuron predictions are shown across models architectures, model layers and neurons. **D.** Pairwise comparison of single-neuron explained variance between layers of one example deep neural network model (same as in Figure 4) and baseline linear encoding model using muscle spindle inputs for NHP S (CN, left column; N = 47 neurons) and NHP H (S1, right column; N = 27 neurons) during active movements. Each row represents the comparison performed for the specific layer of the network. Statistical significance was computed with Wilcoxon signed-rank test (*** = *p <* 0.0001; ** = *p <* 0.001; * = *p <* 0.05). Deep layers of task-driven models are the ones that significantly outperform linear models. **E.** Same as in panel D but for passive trials.

**Supplementary Figure 5:**
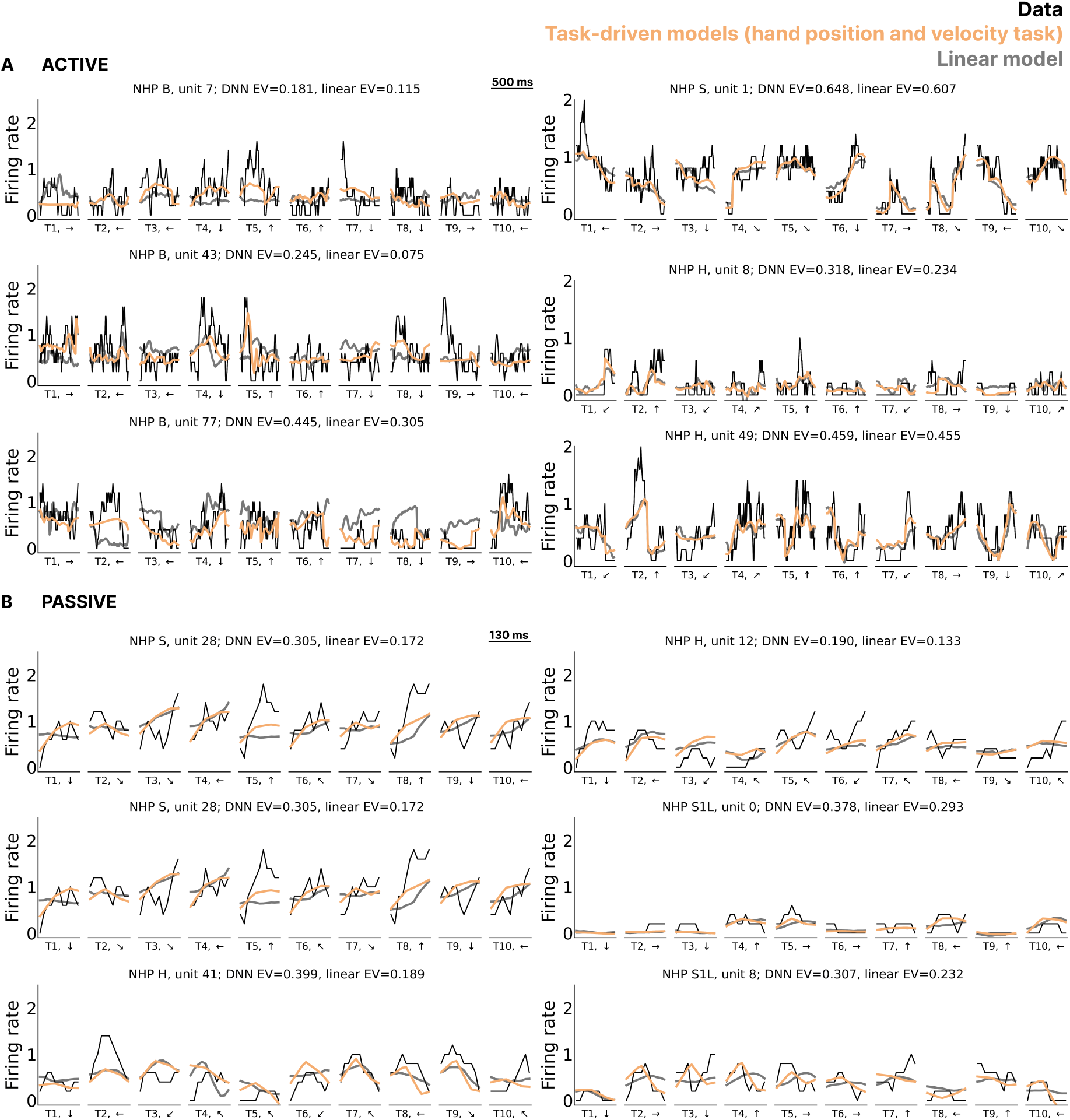
Additional example single-trial predictions, during **A.** active and **B.** passive movement, from the hand position and velocity task (orange) and linear predictions (grey) for different example units in CN and S1. The black lines corresponds to ground truth spike firing rates. Model predictions correspond to the best layer of an example model, the same as in Figure 4. Test performance for task-driven models and linear models is shown. Movement direction is shown for each trial. Task-driven models can capture part of the neural activity that linear models cannot.

**Supplementary Figure 6:**
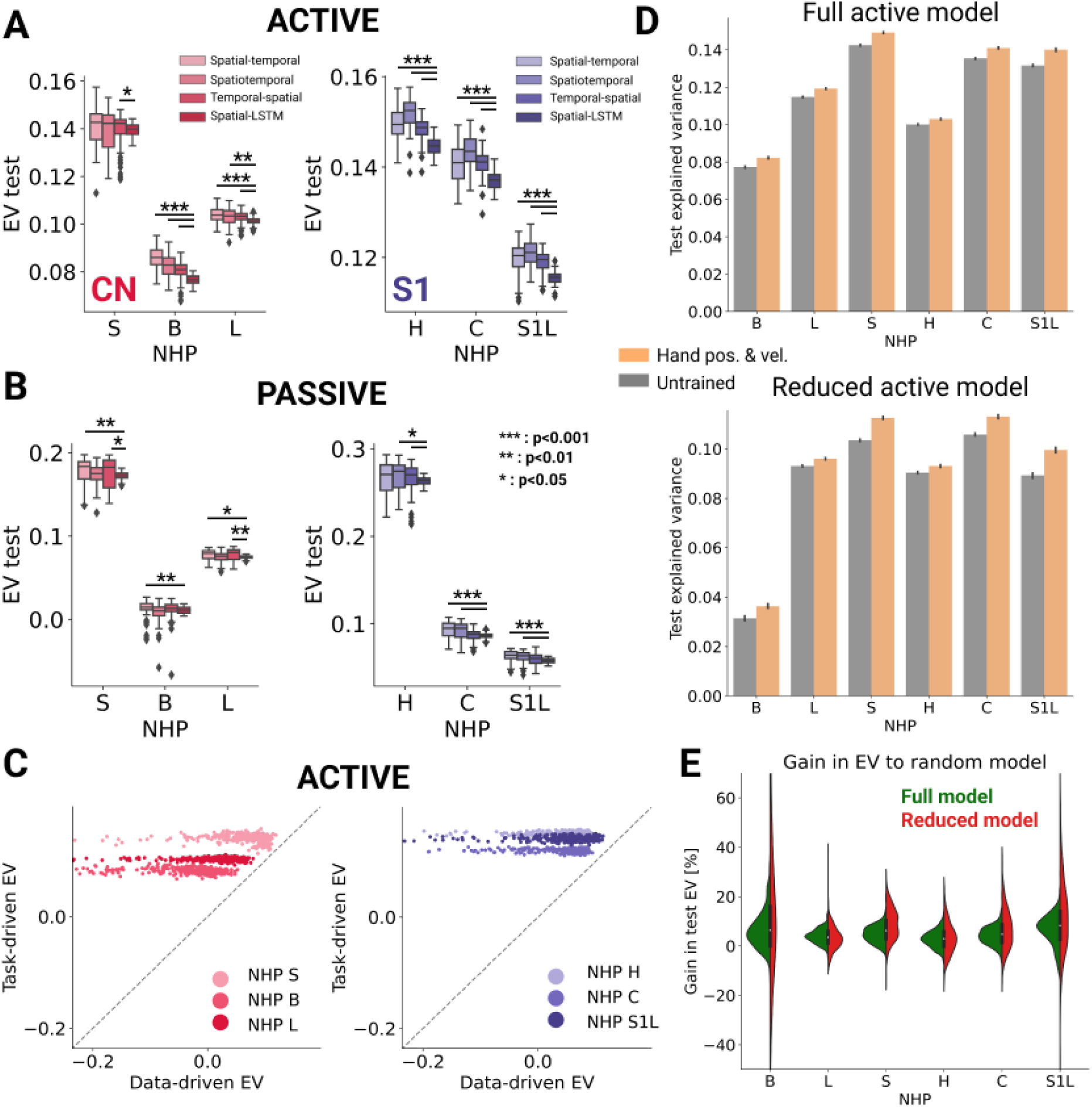
**A.** Distribution of test explained variance between architecture type (N = 100 models per TCN subtype and N = 50 models per Spatial-LSTM models) trained to regress position and velocity of the end-effector for each NHP (CN: red, S1: purple) during active movements. Statistical significance was computed between all TCNs subtype and the spatial-LSTM architecture performing a Mann-Whitney U rank test with Bonferroni correction (only significant comparisons are displayed). Spatial-LSTM is significantly worse at predicting the neural activity, therefore we focused our analysis only on TCN networks. **B.** Same as panel A but for passive movements. **C.** Pairwise comparison between neural explainability obtained from task-driven neural network models trained on the hand position and velocity task (HP & HV) and data-driven neural network models during active movements, for CN (left) and S1 (right) for *N* = 300 TCN models. The data-driven networks have been obtained by training the neural networks to directly regress the neural activity end-to-end. Task-driven models outperform data-driven ones supporting task-driven modeling as a framework for studying tbe proprioceptive systems. **D.** Test neural explainability for networks (N = 300) trained on the hand position and velocity task (HP & HV) and of untrained networks, for each NHP. Top: Full model, i.e. when using all data for the active movements. Bottom: Reduced model, i.e. when active predictions are performed by matching data size to that of passive movement data. In both cases, task-driven networks outperform untrained models thereby discarding the amount of data as a possible cause. **E.** Distribution of the gain explained variance of task-driven networks have with respect to the random models for the full active data and control reduced model.

**Supplementary Figure 7:**
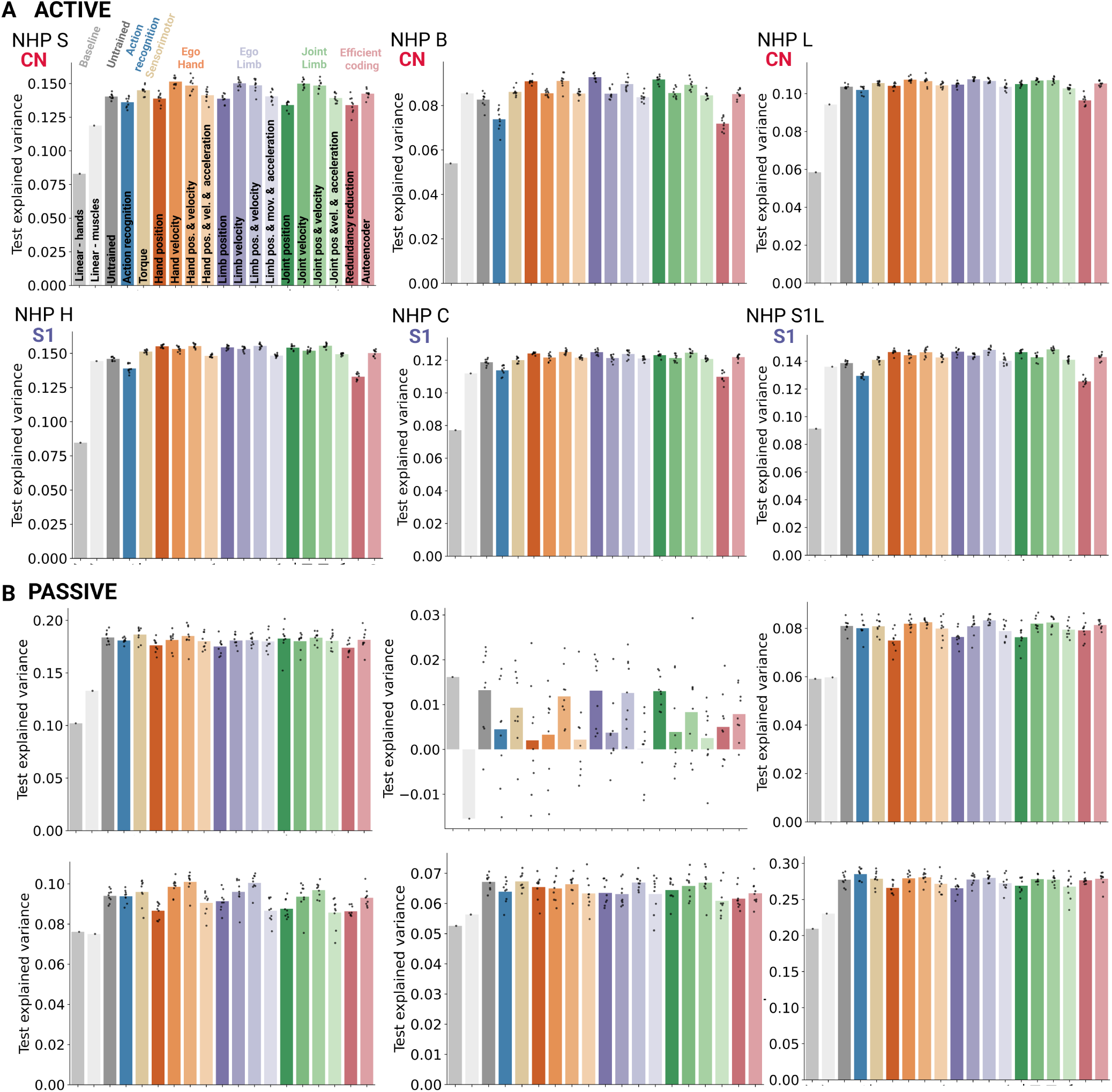
**A.** Explained variance across tasks for all NHPS, as in Figure 5. **B.** Same as panel A but for passive movements, with the same subfigure order per NHP. Note the poor predictions for NHP B in the passive condition.

**Supplementary Figure 8:**
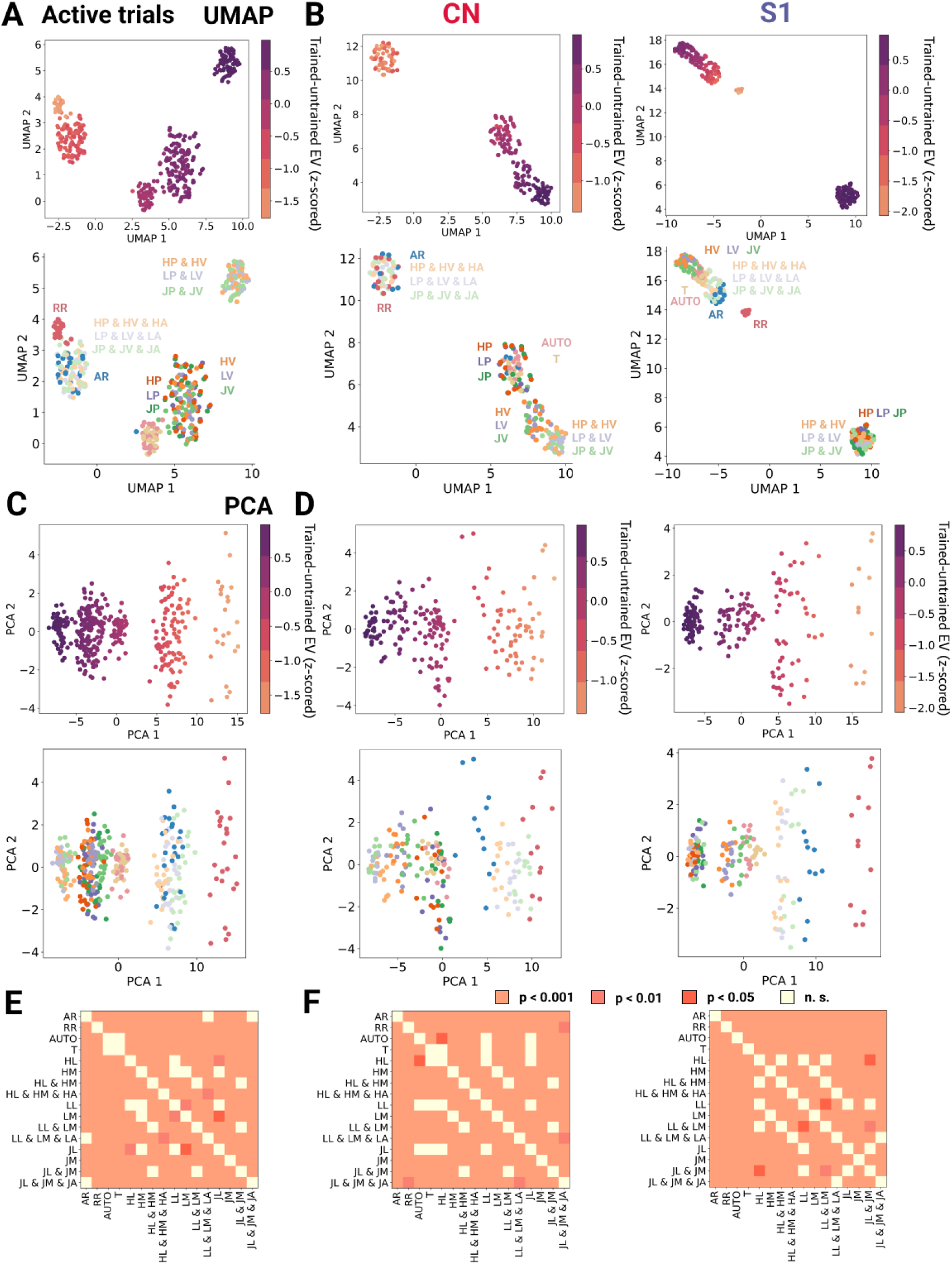
**A.** Low-dimensional UMAP embedding space of the difference in neural explainability between task-driven and untrained networks for active trials. Groups of network belonging to the same task were embedded using the test explained variance by randomly sampling 4 groups of network architectures (N=73 models per group) for each task (N= 16 tasks). The neural explainability was z-scored for each primate and subsequently combined. Top: we color coded each point based on the corresponding average neural explainability. Bottom: we color coded each point according to the task identity. **B.** Same as panel A but computed independently per each brain area (left: CN, right: S1). **C** Same as panel A but the difference is projected along the two first principal components. The same clusters emerge also in the PC space which is a free-parameter approach showing the robustness of the results visualized in the UMAP space **D.** The same as in B but using PCA. **E.** Wilcoxon signed-rank test with Bonferroni correction comparing the gain in explained variance between trained and untrained networks across all tasks. **F.** The same as in C but independently per brain area: left CN, right S1.

**Supplementary Figure 9:**
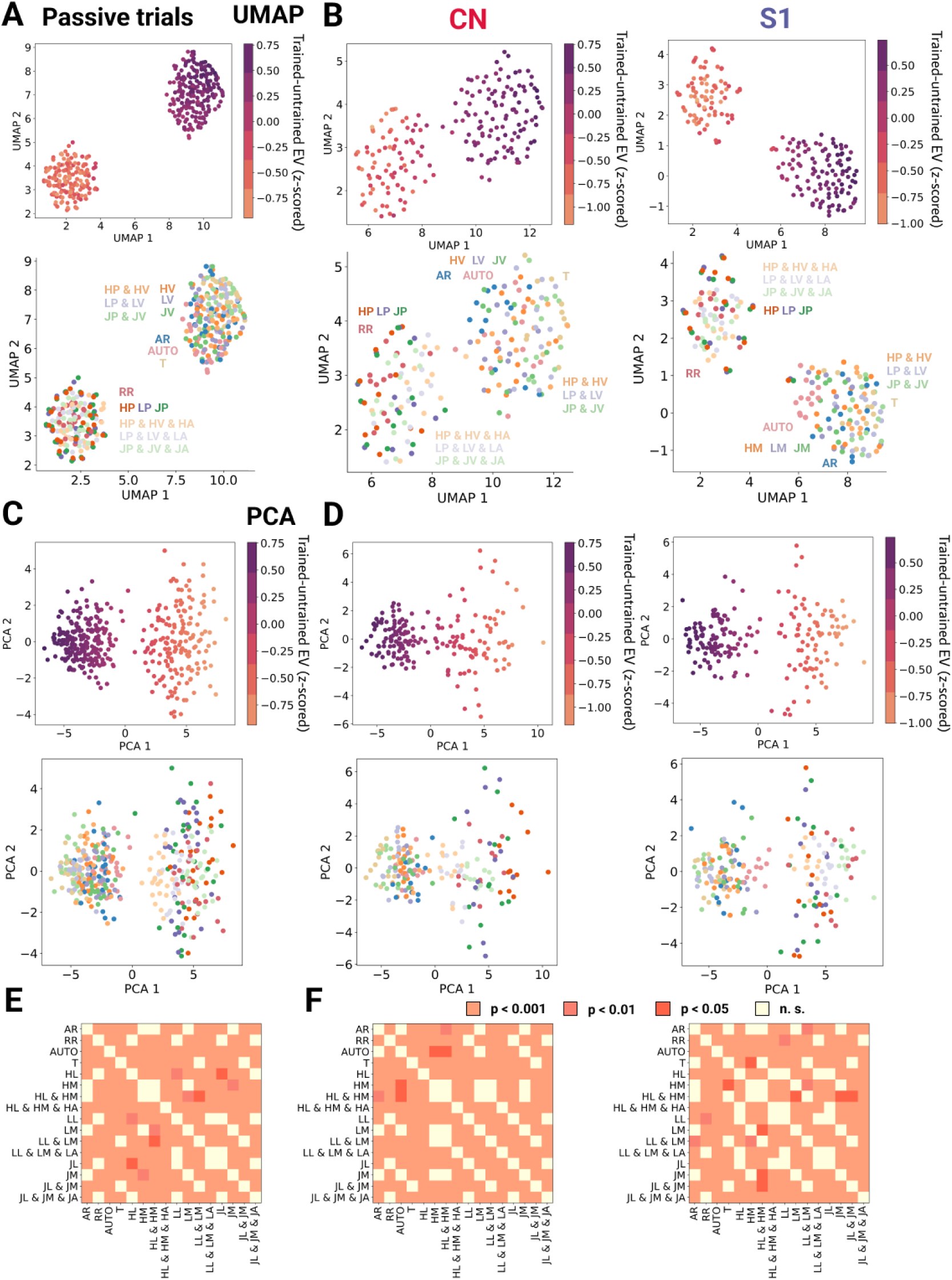
**A.** Low-dimensional UMAP embedding space of the difference in neural explainability between task-driven and untrained networks for passive trials. Groups of network belonging to the same task were embedded using the test explained variance by randomly sampling 4 groups of network architectures (N=73 models per group) for each task (N= 16 tasks). The neural explainability was z-scored for each primate and subsequently combined. Top: we color coded each point based on the corresponding average neural explainability. Bottom: we color coded each point according to the task identity. **B.** Same as panel A but computed independently per each brain area (left: CN, right: S1). **C** Same as panel A but the difference is projected along the two first principal components. The same clusters emerge also in the PC space which is a free-parameter approach showing the robustness of the results visualized in the UMAP space **D.** The same as in B but using PCA. **E.** Wilcoxon signed-rank test with Bonferroni correction comparing the gain in explained variance between trained and untrained networks across all tasks. **F.** The same as in C but independently per brain area: left CN, and right S1.

**Supplementary Figure 10:**
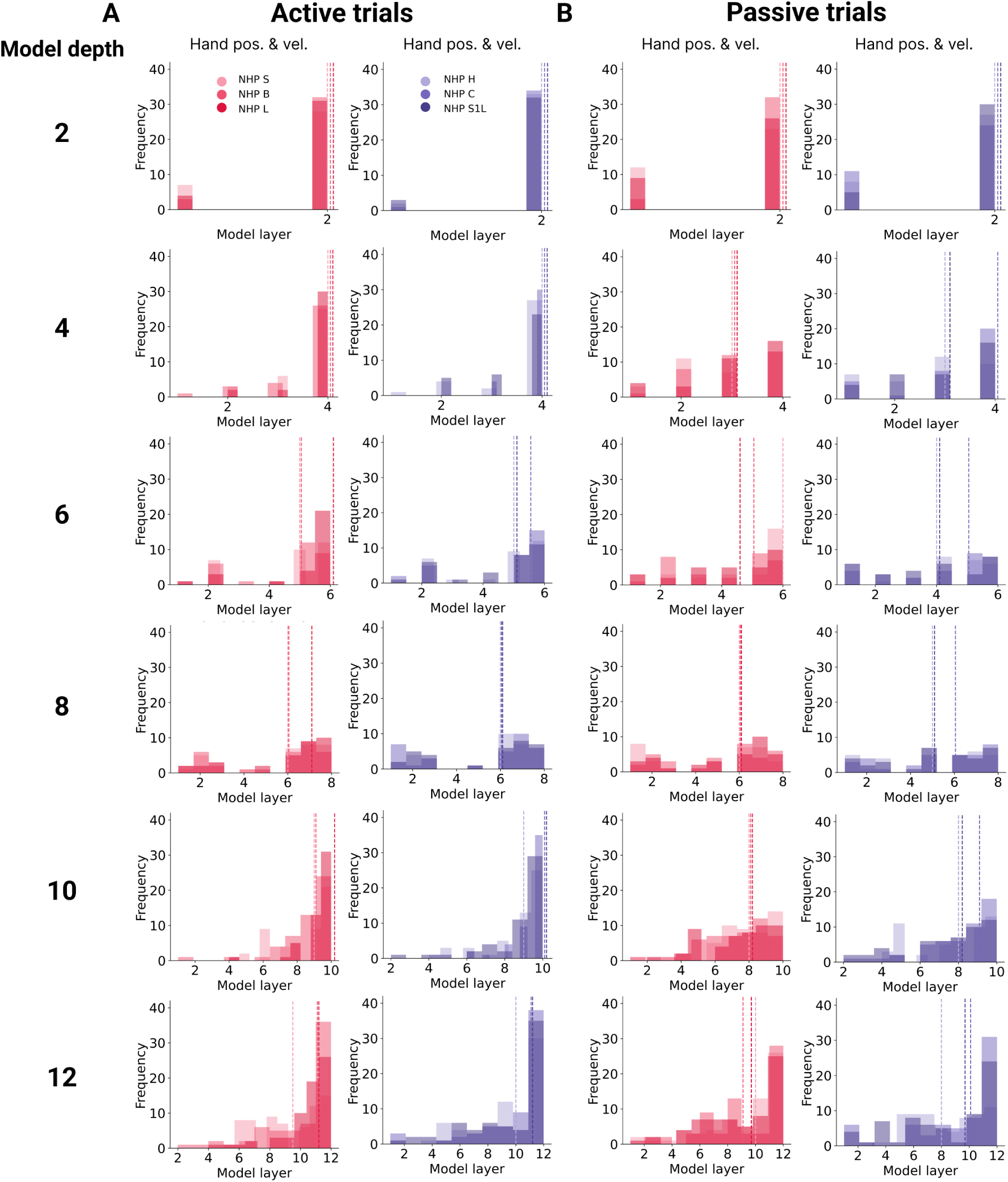
**A.** Empirical distribution of the best predicting layers across all models (N = 350) trained on the hand position and velocity task for CN (left) and S1 (right) for different depth. The dashed line represents the median of the distribution of each NHP. Both CN and S1 NHPs are best predicted by last layer of the networks. As those layers are the ones which more kinematic-tuned features, it suggests that neural activity in both brain areas are tuned to kinematic information without the emergence of any clear hierarchy. **B.** Same as panel A but for passive movements.

**Supplementary Figure 11:**
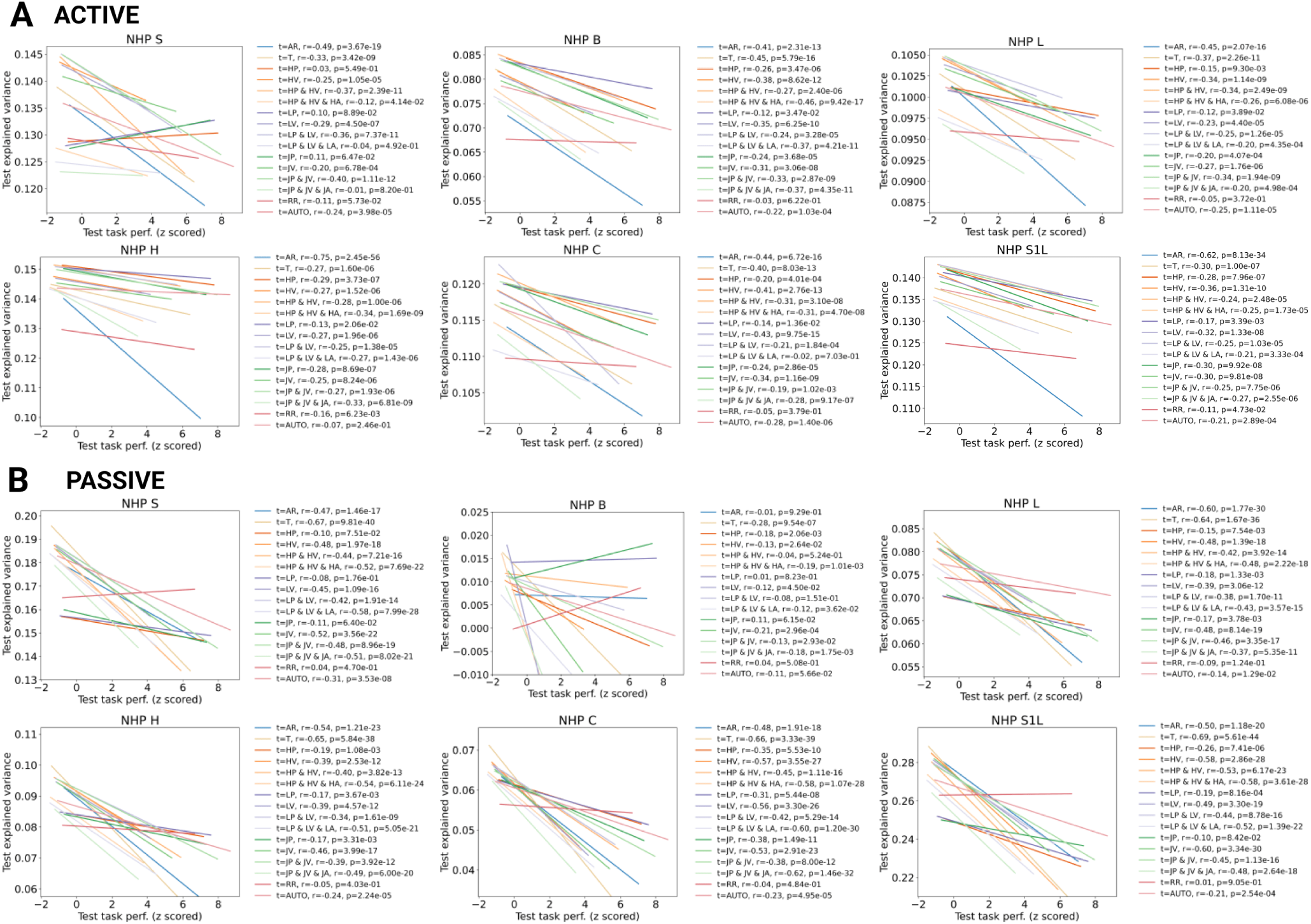
**A.** Linear fits and correlation coefficients showing the relationship between test explained variance of neural predictions, and z-scored networks’ task performance for all tasks. Pearson’s r correlation coefficients and p-values are shown for each NHPs (CN m ∈ [B, L, S], top row; S1 m ∈ [H, C, S1L], bottom row) and each task which is shown with a different colors. For each task, the linear fit was computed using 300 TCN architectures (100 per TCN subtype: spatiotemporal, temporal-spatial, spatial-temporal). Correlation is significant across NHPs and across tasks underlying the robustness of task-optimization for achieving better neural network representation to predict the neural activity. **B.** Same as panel A but for passive movements.

**Supplementary Figure 12:**
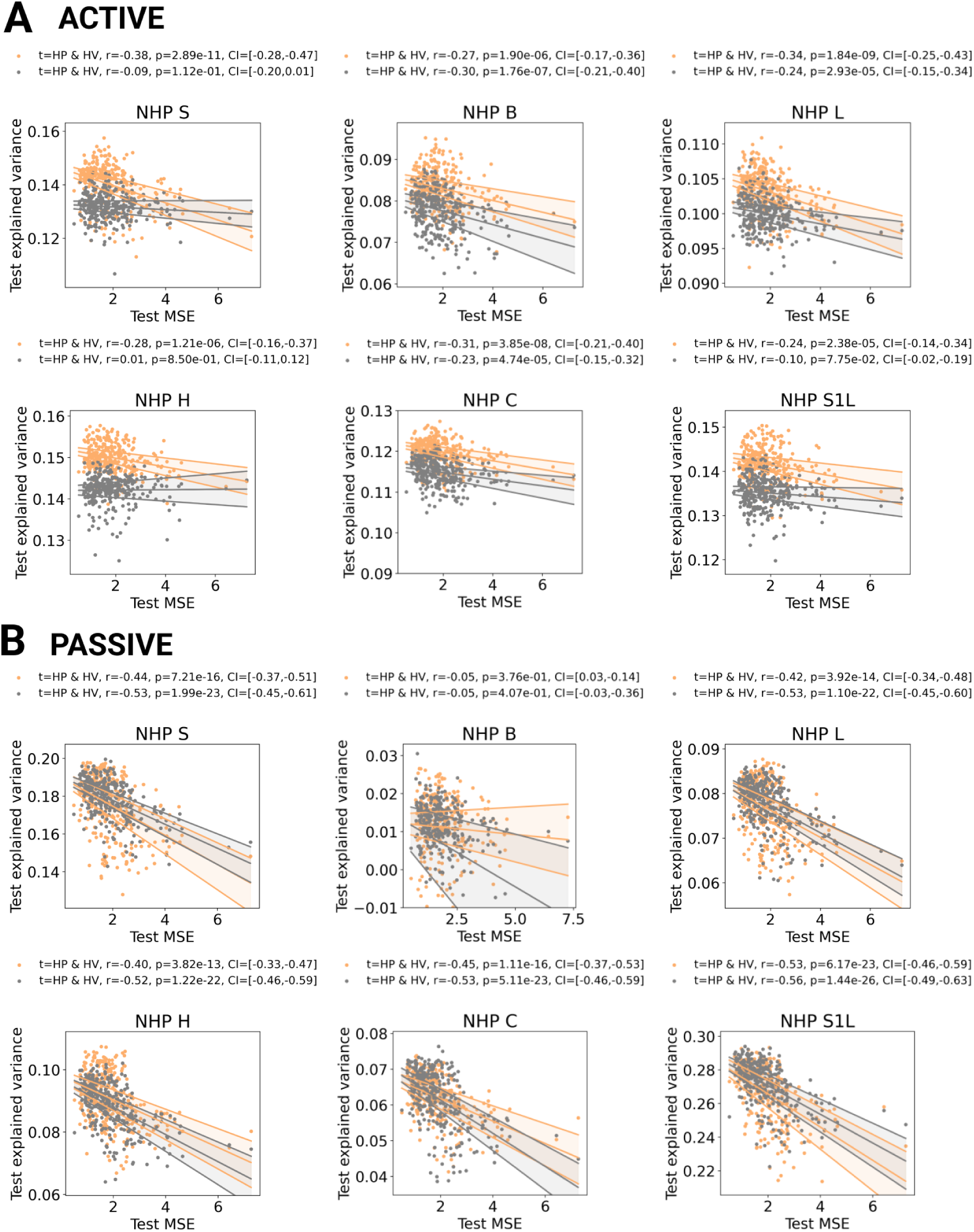
**A.** Scatter plot of the relationship between test task performance and neural explainability for the hand position and velocity task (orange) and random, untrained models (gray). Explained variance of random models is ordered based on the task performance of the corresponding task-driven network architecture. Pearson’s *r* correlation coefficients and p-values are shown for each NHPs (CN m ∈ [B, L, S], top row; S1 m ∈ [H, C, S1L], bottom row). Confidence interval (shaded area) computed with bootstrap by resampling 1000 times. Each data point represents the neural explainability of a single model. In total, N=300 TCN architectures (100 per TCN subtype: spatiotemporal, temporal-spatial, spatial-temporal) are shown. Lines indicate linear fits for each NHPs. **B.** Same as panel A but for passive movements. Correlation of task-trained network outperform the untrained one for active data but not for passive one suggesting the neural activity is differently processed during the two conditions: neural activity might top-down modulated with motor efference copy during goal-directed movements whereas it is likely to be absent during passive movements.

**Supplementary Figure 13:**
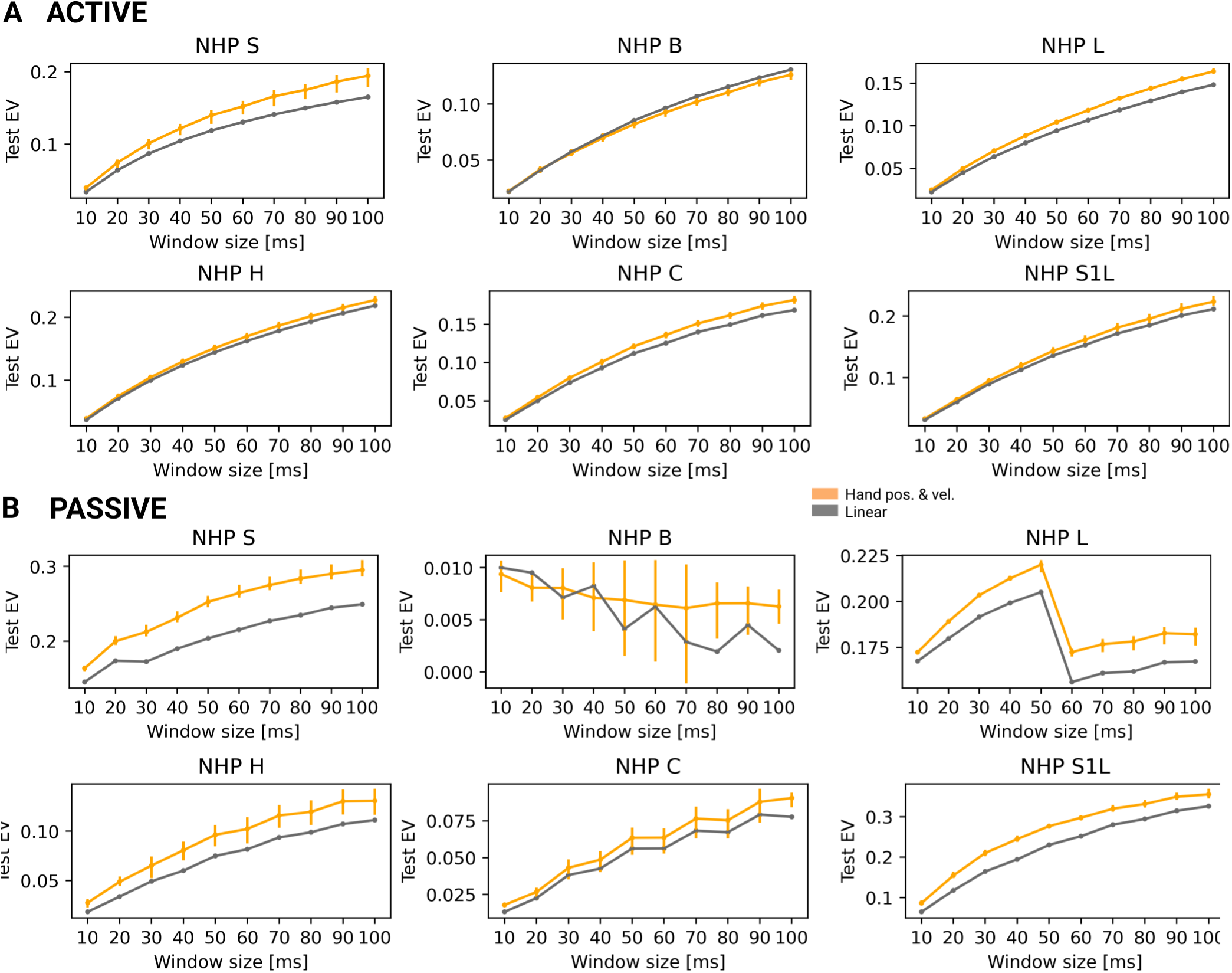
**A.** Neural explainability for different sizes of window used to compute spike rates, for three example neural network models (one per TCN subtype, including model of Fig.4) of the hand position and velocity task (color) and for linear models (gray). Error bars represent 95% confidence intervals. **B.** Same as panel A for passive predictions. Task-driven models typically outperform linear models independent of the window size both in CN and S1.

**Supplementary Figure 14:**
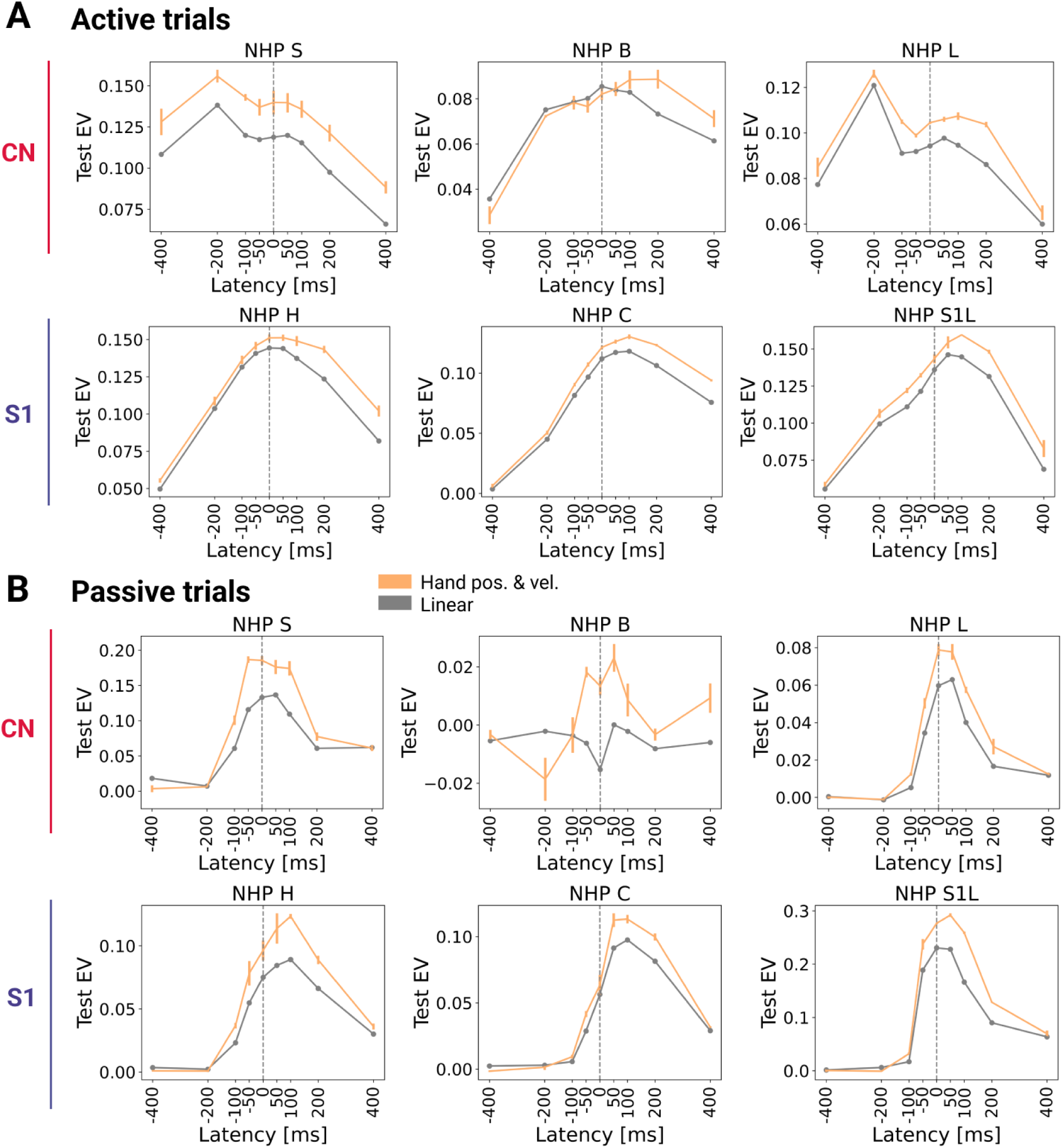
**A.** Neural explainability for different latency shifts, for three example neural network models (one per TCN subtype, including model of Fig.4) of the hand position and velocity task. Error bars represent 95% confidence intervals. **B.** Same as panel A for passive predictions. Task-driven models typically outperform linear models for a wide range o latencies around 0 both in CN and S1.

## Notes

### Competing Interest Statement

The authors have declared no competing interest.

## References

1. Daniel LK Yamins and James J DiCarlo. Using goal-driven deep learning models to understand sensory cortex. Nature neuroscience, 19(3):356, 2016.

2. Blake A Richards, Timothy P Lillicrap, Philippe Beaudoin, Yoshua Bengio, Rafal Bogacz, Amelia Christensen, Claudia Clopath, Rui Ponte Costa, Archy de Berker, Surya Ganguli, et al. A deep learning framework for neuroscience. Nature Neuroscience, 22(11):1761–1770, 2019.

3. Sébastien B Hausmann, Alessandro Marin Vargas, Alexander Mathis, and Mackenzie W Mathis. Measuring and modeling the motor system with machine learning. Current opinion in neurobiology, 70:11–23, 2021.

4. Rosa Cao and Daniel Yamins. Explanatory models in neuroscience: Part 2–constraint-based intelligibility. *arXiv preprint arXiv:2104.01489*, 2021.

5. Daniel LK Yamins, Ha Hong, Charles F Cadieu, Ethan A Solomon, Darren Seibert, and James J DiCarlo. Performance-optimized hierarchical models predict neural responses in higher visual cortex. Proceedings of the national academy of sciences, 111(23):8619–8624, 2014.

6. Alexander JE Kell, Daniel LK Yamins, Erica N Shook, Sam V Norman-Haignere, and Josh H McDermott. A task-optimized neural network replicates human auditory behavior, predicts brain responses, and reveals a cortical processing hierarchy. Neuron, 98(3):630–644, 2018.

7. Martin Haesemeyer, Alexander F Schier, and Florian Engert. Convergent temperature representations in artificial and biological neural networks. Neuron, 103(6):1123–1134, 2019.

8. Jonathan A Michaels, Stefan Schaffelhofer, Andres Agudelo-Toro, and Hansjörg Scherberger. A goal-driven modular neural network predicts parietofrontal neural dynamics during grasping. Proceedings of the national academy of sciences, 117(50):32124–32135, 2020.

9. Chengxu Zhuang, Siming Yan, Aran Nayebi, Martin Schrimpf, Michael C Frank, James J DiCarlo, and Daniel LK Yamins. Unsupervised neural network models of the ventral visual stream. Proceedings of the National Academy of Sciences, 118(3), 2021.

10. Aran Nayebi, Javier Sagastuy-Brena, Daniel M Bear, Kohitij Kar, Jonas Kubilius, Surya Ganguli, David Sussillo, James J DiCarlo, and Daniel LK Yamins. Goal-driven recurrent neural network models of the ventral visual stream. bioRxiv, 2021.

11. Ariel Goldstein, Zaid Zada, Eliav Buchnik, Mariano Schain, Amy Price, Bobbi Aubrey, Samuel A Nastase, Amir Feder, Dotan Emanuel, Alon Cohen, et al. Shared computational principles for language processing in humans and deep language models. Nature neuroscience, 25(3):369–380, 2022.

12. Aran Nayebi, Alexander Attinger, Malcolm Campbell, Kiah Hardcastle, Isabel Low, Caitlin S Mallory, Gabriel Mel, Ben Sorscher, Alex H Williams, Surya Ganguli, et al. Explaining heterogeneity in medial entorhinal cortex with task-driven neural networks. Advances in Neural Information Processing Systems, 34:12167–12179, 2021.

13. Santiago A Cadena, Konstantin F Willeke, Kelli Restivo, George Denfield, Fabian H Sinz, Matthias Bethge, Andreas S Tolias, and Alexander S Ecker. Diverse task-driven modeling of macaque v4 reveals functional specialization towards semantic tasks. bioRxiv, pages 2022–05, 2022.

14. Pouya Bashivan, Kohitij Kar, and James J DiCarlo. Neural population control via deep image synthesis. Science, 364 (6439):eaav9436, 2019.

15. Kai J Sandbrink, Pranav Mamidanna, Claudio Michaelis, Matthias Bethge, Mackenzie W Mathis, and Alexander Mathis. Contrasting action and posture coding with hierarchical deep neural network models of proprioception. Elife, 12:e81499, 2023.

16. Kyle P Blum, Max D Grogan, Yufei Wu, Alex J Harston, Lee E Miller, and Aldo A Faisal. Predicting proprioceptive cortical anatomy and neural coding with topographic autoencoders. bioRxiv, 2021.

17. H Charlton Bastian. The “muscular sense”; its nature and cortical localisation. Brain, 10(1):1–89, 1887.

18. Stephen H Scott. Optimal feedback control and the neural basis of volitional motor control. Nature Reviews Neuroscience, 5(7):532–545, 2004.

19. Uwe Proske and Simon C Gandevia. The proprioceptive senses: their roles in signaling body shape, body position and movement, and muscle force. Physiological reviews, 2012.

20. Benoit P Delhaye, Katie H Long, and Sliman J Bensmaia. Neural basis of touch and proprioception in primate cortex. Comprehensive Physiology, 8(4):1575, 2018.

21. Arthur Prochazka and Monica Gorassini. Models of ensemble firing of muscle spindle afferents recorded during normal locomotion in cats. The Journal of physiology, 507(1):277– 291, 1998.

22. Milana P. Mileusnic, Ian E. Brown, Ning Lan, and Gerald E. Loeb. Mathematical models of proprioceptors. i. control and transduction in the muscle spindle. Journal of Neurophysiology, 96(4):1772–1788, Oct 2006. ISSN 0022-3077. doi: 10.1152/jn.00868.2005.

23. Michael Dimitriou and Benoni B Edin. Discharges in human muscle spindle afferents during a key-pressing task. The Journal of physiology, 586(22):5455–5470, 2008.

24. Kyle P Blum, Kenneth S Campbell, Brian C Horslen, Paul Nardelli, Stephen N Housley, Timothy C Cope, and Lena H Ting. Diverse and complex muscle spindle afferent firing properties emerge from multiscale muscle mechanics. Elife, 9:e55177, 2020.

25. Michael Dimitriou. Human muscle spindles are wired to function as controllable signal-processing devices. Elife, 11: e78091, 2022.

26. P. Andersen, J. C. Eccles, T. Oshima, and R. F. Schmidt. Mechanisms of synaptic transmission in the cuneate nucleus. Journal of Neurophysiology, 27(6):1096–1116, Nov 1964. ISSN 0022-3077, 1522-1598. doi: 10.1152/jn.1964.27.6.1096.

27. P. Andersen, J. C. Eccles, R. F. Schmidt, and T. Yokota. Identification of relay cells and interneurons in the cuneate nucleus. Journal of Neurophysiology, 27(6):1080–1095, Nov 1964. ISSN 0022-3077, 1522-1598. doi: 10.1152/jn.1964.27.6.1080.

28. Marion C. Smith and Patricia Deacon. Topographical anatomy of the posterior columns of the spinal cord in man: the long ascending fibres. Brain, 107(3):671–698, 1984. ISSN 0006-8950, 1460-2156. doi: 10.1093/brain/107.3.671.

29. Leah A Krubitzer and Jon H Kaas. The somatosensory thalamus of monkeys: cortical connections and a redefinition of nuclei in marmosets. Journal of Comparative Neurology, 319(1):123–140, 1992.

30. Joseph T Francis, Shaohua Xu, and John K Chapin. Proprioceptive and cutaneous representations in the rat ventral posterolateral thalamus. Journal of neurophysiology, 99(5): 2291–2304, 2008.

31. MJ Prud’Homme and John F Kalaska. Proprioceptive activity in primate primary somatosensory cortex during active arm reaching movements. Journal of neurophysiology, 72 (5):2280–2301, 1994.

32. Alastair J. Loutit, Richard M. Vickery, and Jason R. Potas. Functional organization and connectivity of the dorsal column nuclei complex reveals a sensorimotor integration and distribution hub. Journal of Comparative Neurology, page cne.24942, Jun 2020. ISSN 0021-9967, 1096-9861. doi: 10.1002/cne.24942.

33. Rafael V Bretas, Miki Taoka, Hiroaki Suzuki, and Atsushi Iriki. Secondary somatosensory cortex of primates: beyond body maps, toward conscious self-in-the-world maps. Experimental Brain Research, 238(2):259–272, 2020.

34. Christoph Fromm and Edward V Evarts. Pyramidal tract neurons in somatosensory cortex: central and peripheral inputs during voluntary movement. Brain research, 238(1): 186–191, 1982.

35. Brian M London and Lee E Miller. Responses of somatosensory area 2 neurons to actively and passively generated limb movements. Journal of neurophysiology, 109(6):1505–1513, 2013.

36. Christopher Versteeg, Joshua M Rosenow, Sliman J Bensmaia, and Lee E Miller. Encoding of limb state by single neurons in the cuneate nucleus of awake monkeys. Journal of Neurophysiology, 2021.

37. Volker Dietz. Proprioception and locomotor disorders. Nature Reviews Neuroscience, 3(10):781–790, 2002.

38. Jonathan S Tsay, Hyosub Kim, Adrian M Haith, and Richard B Ivry. Understanding implicit sensorimotor adaptation as a process of proprioceptive re-alignment. eLife, 11: e76639, aug 2022. ISSN 2050-084X. doi: 10.7554/eLife.76639.

39. Horace B Barlow et al. Possible principles underlying the transformation of sensory messages. Sensory communication, 1(01), 1961.

40. Eero P Simoncelli and Bruno A Olshausen. Natural image statistics and neural representation. Annual review of neuroscience, 24(1):1193–1216, 2001.

41. Raeed H Chowdhury, Joshua I Glaser, and Lee E Miller. Area 2 of primary somatosensory cortex encodes kinematics of the whole arm. Elife, 9:e48198, 2020.

42. John C Rothwell. Control of human voluntary movement. Springer Science & Business Media, 2012.

43. Jure Zbontar, Li Jing, Ishan Misra, Yann LeCun, and Stéphane Deny. Barlow twins: Self-supervised learning via redundancy reduction. In International Conference on Machine Learning, pages 12310–12320. PMLR, 2021.

44. Alexander Mathis, Pranav Mamidanna, Kevin M Cury, Taiga Abe, Venkatesh N Murthy, Mackenzie Weygandt Mathis, and Matthias Bethge. Deeplabcut: markerless pose estimation of user-defined body parts with deep learning. Nature neuroscience, 21(9):1281–1289, 2018.

45. Alice Lucas, Tucker Tomlinson, Neda Rohani, Raeed Chowdhury, Sara A Solla, Aggelos K Katsaggelos, and Lee E Miller. Neural networks for modeling neural spiking in s1 cortex. Frontiers in systems neuroscience, 13:13, 2019.

46. James M Goodman, Gregg A Tabot, Alex S Lee, Aneesha K Suresh, Alexander T Rajan, Nicholas G Hatsopoulos, and Sliman Bensmaia. Postural representations of the hand in the primate sensorimotor cortex. Neuron, 104(5): 1000–1009, 2019.

47. Mackenzie Weygandt Mathis, Alexander Mathis, and Naoshige Uchida. Somatosensory cortex plays an essential role in forelimb motor adaptation in mice. Neuron, 93(6): 1493–1503, 2017.

48. Jose M Carmena, Mikhail A Lebedev, Roy E Crist, Joseph E O’Doherty, David M Santucci, Dragan F Dimitrov, Parag G Patil, Craig S Henriquez, and Miguel A L Nicolelis. Learning to control a brain–machine interface for reaching and grasping by primates. PLoS biology, 1(2):e42, 2003.

49. Christopher Versteeg, Raeed H Chowdhury, and Lee E Miller. Cuneate nucleus: The somatosensory gateway to the brain. Current Opinion in Physiology, 20:206–215, 2021.

50. Daniel M Wolpert, Susan J Goodbody, and Masud Husain. Maintaining internal representations: the role of the human superior parietal lobe. Nature neuroscience, 1(6):529–533, 1998.

51. Uwe Proske, David L Morgan, and JE Gregory. Muscle history dependence of responses to stretch of primary and secondary endings of cat soleus muscle spindles. The Journal of Physiology, 445(1):81–95, 1992.

52. Peter BC Matthews. Muscle spindles: their messages and their fusimotor supply. Comprehensive Physiology, pages 189–228, 2011.

53. Kyle P Blum, Boris Lamotte D’Incamps, Daniel Zytnicki, and Lena H Ting. Force encoding in muscle spindles during stretch of passive muscle. PLoS computational biology, 13 (9):e1005767, 2017.

54. Vaughan G Macefield and Thomas P Knellwolf. Functional properties of human muscle spindles. Journal of neurophysiology, 120(2):452–467, 2018.

55. Edward V Evarts. Relation of pyramidal tract activity to force exerted during voluntary movement. Journal of neurophysiology, 31(1):14–27, 1968.

56. Eberhard E Fetz and Paul D Cheney. Postspike facilitation of forelimb muscle activity by primate corticomotoneuronal cells. Journal of neurophysiology, 44(4):751– 772, 1980.

57. Apostolos P Georgopoulos, John F Kalaska, Roberto Caminiti, and Joe T Massey. On the relations between the direction of two-dimensional arm movements and cell discharge in primate motor cortex. Journal of Neuroscience, 2 (11):1527–1537, 1982.

58. Apostolos P Georgopoulos, Ronald E Kettner, and Andrew B Schwartz. Primate motor cortex and free arm movements to visual targets in three-dimensional space. ii. coding of the direction of movement by a neuronal population. Journal of Neuroscience, 8(8):2928–2937, 1988.

59. John F Kalaska. From intention to action: motor cortex and the control of reaching movements. Progress in Motor Control: A Multidisciplinary Perspective, pages 139–178, 2009.

60. Gianfranco Bosco, Richard E Poppele, and John Eian. Reference frames for spinal proprioception: limb endpoint based or joint-level based? Journal of Neurophysiology, 83 (5):2931–2945, 2000.

61. Erich von Holst and H Mittelstaedt. The reafference principle (r. martin, trans.). the behavioral physiology of animals and man: The collected papers of erich von holst, 1973.

62. Roger Wolcott Sperry. Neural basis of the spontaneous optokinetic response produced by visual inversion. Journal of comparative and physiological psychology, 43(6):482, 1950.

63. Tatsuya Umeda, Tadashi Isa, and Yukio Nishimura. The somatosensory cortex receives information about motor output. Science advances, 5(7):eaaw5388, 2019.

64. Fang Cui, Dan Arnstein, Rajat Mani Thomas, Natasha M Maurits, Christian Keysers, and Valeria Gazzola. Functional magnetic resonance imaging connectivity analyses reveal efference-copy to primary somatosensory area, ba2. PLoS One, 9(1):e84367, 2014.

65. John Kalambogias and Yutaka Yoshida. Converging integration between ascending proprioceptive inputs and the corticospinal tract motor circuit underlying skilled movement control. Current opinion in physiology, 19:187–193, 2021.

66. Alastair J. Loutit, Richard M. Vickery, and Jason R. Potas. Functional organization and connectivity of the dorsal column nuclei complex reveals a sensorimotor integration and distribution hub. Journal of Comparative Neurology, 529(1): 187–220, 2021.

67. Jorge Mariño, Luis Martinez, and Antonio Canedo. Sensorimotor integration at the dorsal column nuclei. Physiology, 14(6):231–237, 1999.

68. Arthur Prochazka, M. Hulliger, P. Zangger, and K. Appenteng. ‘fusimotor set’: new evidence for *α*-independent control of *γ*-motoneurones during movement in the awake cat. Brain research, 339(1):136–140, 1985.

69. Joachim Confais, Geehee Kim, Saeka Tomatsu, Tomohiko Takei, and Kazuhiko Seki. Nerve-specific input modulation to spinal neurons during a motor task in the monkey. Journal of Neuroscience, 37(10):2612–2626, 2017.

70. Sung Soo Kim, Manuel Gomez-Ramirez, Pramodsingh H Thakur, and Steven S Hsiao. Multimodal interactions between proprioceptive and cutaneous signals in primary somatosensory cortex. Neuron, 86(2):555–566, 2015.

71. JF Kalaska, DAD Cohen, M Prud’Homme, and ML Hyde. Parietal area 5 neuronal activity encodes movement kinematics, not movement dynamics. Experimental brain research, 80(2):351–364, 1990.

72. Jeffrey Padberg, Dylan F Cooke, Christina M Cerkevich, Jon H Kaas, and Leah Krubitzer. Cortical connections of area 2 and posterior parietal area 5 in macaque monkeys. Journal of Comparative Neurology, 527(3):718–737, 2019.

73. Huawei Wang, Vittorio Caggiano, Guillaume Durandau, Massimo Sartori, and Vikash Kumar. Myosim: Fast and physiologically realistic mujoco models for musculoskeletal and exoskeletal studies. In 2022 International Conference on Robotics and Automation (ICRA), pages 8104–8111. IEEE, 2022.

74. Alberto Silvio Chiappa, Alessandro Marin Vargas, Ann Zixiang Huang, and Alexander Mathis. Latent exploration for reinforcement learning. arXiv preprint arXiv:2305.20065, 2023.

75. Pierre Schumacher, Daniel Häufle, Dieter Büchler, Syn Schmitt, and Georg Martius. Dep-rl: Embodied exploration for reinforcement learning in overactuated and musculoskeletal systems. arXiv preprint arXiv:2206.00484, 2022.

76. Timothy P Lillicrap and Stephen H Scott. Preference distributions of primary motor cortex neurons reflect control solutions optimized for limb biomechanics. Neuron, 77(1):168– 179, 2013.

77. Maria Knikou. The h-reflex as a probe: pathways and pitfalls. Journal of neuroscience methods, 171(1):1–12, 2008.

78. Ariel J Levine, Kathryn A Lewallen, and Samuel L Pfaff. Spatial organization of cortical and spinal neurons controlling motor behavior. Current opinion in neurobiology, 22(5): 812–821, 2012.

79. Alberto Silvio Chiappa, Alessandro Marin Vargas, and Alexander Mathis. Dmap: a distributed morphological attention policy for learning to locomote with a changing body. Advances in Neural Information Processing Systems, 35: 37214–37227, 2022.

80. Aran Nayebi, Daniel Bear, Jonas Kubilius, Kohitij Kar, Surya Ganguli, David Sussillo, James J DiCarlo, and Daniel L Yamins. Task-driven convolutional recurrent models of the visual system. Advances in neural information processing systems, 31, 2018.

81. Jonas Kubilius, Martin Schrimpf, Kohitij Kar, Rishi Rajalingham, Ha Hong, Najib Majaj, Elias Issa, Pouya Bashivan, Jonathan Prescott-Roy, Kailyn Schmidt, et al. Brain-like object recognition with high-performing shallow recurrent anns. Advances in neural information processing systems, 32, 2019.

82. Aneesha K Suresh, Jeremy E Winberry, Christopher Versteeg, Raeed Chowdhury, Tucker Tomlinson, Joshua M Rosenow, Lee E Miller, and Sliman J Bensmaia. Methodological considerations for a chronic neural interface with the cuneate nucleus of macaques. Journal of Neurophysiology, 118(6):3271–3281, 2017.

83. Tanmay Nath, Alexander Mathis, An Chi Chen, Amir Patel, Matthias Bethge, and Mackenzie Weygandt Mathis. Using deeplabcut for 3d markerless pose estimation across species and behaviors. Nature protocols, 14(7):2152–2176, 2019.

84. Sherwin S Chan and Daniel W Moran. Computational model of a primate arm: from hand position to joint angles, joint torques and muscle forces. Journal of neural engineering, 3(4):327, 2006.

85. Scott L Delp, Frank C Anderson, Allison S Arnold, Peter Loan, Ayman Habib, Chand T John, Eran Guendelman, and Darryl G Thelen. Opensim: open-source software to create and analyze dynamic simulations of movement. IEEE transactions on biomedical engineering, 54(11):1940–1950, 2007.

86. Ajay Seth, Michael Sherman, Jeffrey A Reinbolt, and Scott L Delp. Opensim: a musculoskeletal modeling and simulation framework for in silico investigations and exchange. Procedia Iutam, 2:212–232, 2011.

87. Ben H Williams, Marc Toussaint, and Amos J Storkey. Extracting motion primitives from natural handwriting data. In International Conference on Artificial Neural Networks, pages 634–643. Springer, 2006.

88. Arthur Asuncion and David Newman. Uci machine learning repository, 2007.

89. Erwin Coumans and Yunfei Bai. Pybullet, a python module for physics simulation for games, robotics and machine learning, 2016.

90. Tuomas Haarnoja, Aurick Zhou, Pieter Abbeel, and Sergey Levine. Soft actor-critic: Off-policy maximum entropy deep reinforcement learning with a stochastic actor. International conference on machine learning, pages 1861–1870, 2018.

91. Leland McInnes, John Healy, and James Melville. Umap: Uniform manifold approximation and projection for dimension reduction. arXiv preprint arXiv:1802.03426, 2018.

